# Generalized inference of noise-corrected state variability in single cells

**DOI:** 10.64898/2026.11.09.765210

**Authors:** Russell B. Hawes, Mercè Garí, Wisam A. Fares, Cecilia J. Fetterolf, Alice Chang, Najwa Labban, Shayla A. Vradenburgh, Lixin Wang, Róża K. Przanowska, Hui Zong, Shayna L. Showalter, Christiane Fuchs, Kevin A. Janes

## Abstract

Single-cell omics methods operate at their detection limit and are less conducive to technical controls than “mini-bulk” approaches, which combine cells of the same type before profiling. To integrate mini-bulk samples with paired split-pool controls more holistically, we built stochprofIC, an algorithm that considers data, noise, and covariates in a distributional-regression framework. We vet stochprofIC with synthetic datasets and compare against an adjusted-variance alternative using mini-bulk data in mouse brains and human breast cancers. The noise–batch corrections provided by stochprofIC enable inference of the single-cell populations comprising the mini-bulk, which are validated repeatedly by RNA FISH in different contexts. We use stochprofIC outputs as input to a systems model of nucleocytoplasmic transport and distinguish heterogeneous trafficking states. Pharmacologic inhibition of a *MAP3K11*-high state inhibits apoptosis in primary breast-cancer organoids, supporting relevance to phenotypes. By deconvolving changes in location and scale, stochprofIC supports mini-bulking when high-sensitivity and reliability profiles are important.

**HIGHLIGHTS:**

- stochprofIC infers noise and covariate effects, then corrects mini-bulk profiles
- stochprofIC catalyzes maximum-likelihood inference of a gene’s single-cell states
- Predicted single-cell expression frequencies agree with RNA FISH estimates
- Corrected mini-bulk data inform systems models and follow-up perturbations

## INTRODUCTION

Single-cell RNA sequencing (scRNA-seq) distinguishes qualitative cell types very effectively, but within a cell type, its coverage of subtler, quantitative regulatory states is less clear.^1,2^ Cell type–state distinctions are particularly blurred in cancer, where measured scRNA-seq profiles skew by the copy-number variations of a tumor and the cycling status of individual cells.^3–6^ Upon correcting for or filtering out these confounders, scRNA-seq has identified recurrent intratumor variabilities in differentiation “state” (i.e., lineage) along with robust transcriptional programs downstream of cell stress, interferons, and hypoxia.^7,8^ However, this list may be partial, as whole tissue–tumor transcriptomes are not fully recovered when paired scRNA-seq data are pseudo-bulked.^9,10^ The recognized detection floor^11^ of scRNA-seq raises the question of whether there are more recurrent cell states to be found.^12^

One pragmatic workaround is mini-bulking, which combines tens to hundreds of similar cells before sample processing. Mini-bulks enhance coverage of low-input proteomics^13^ and enable low-yield sequencing from nascent RNA^14^ or laser-capture microdissected tissue.^15–18^ The increased starting material improves precision^19^ and reduces technical zeros^11^ that arise from losses before sequencing.^16^ Mini-bulk pools as small as 10 cells are sufficient to average out cell cycle-phased transcripts^20^ and improve the validity of distributional assumptions^21^ that might otherwise be questionable for single cells.^22,23^ Larger, input-equivalent “split pools” of the cell type add an empirical measure of technical noise, which has similarly been leveraged for plate-based scRNA-seq^24^ but only recently been considered for droplet-based methods.^25^ An obvious drawback is that mini-bulk profiles are not explicitly single cell, but that is addressable with combined experimental–analytical designs that identify overdispersion^20,26,27^ and reconstruct the most-likely single-cell distributions.^21,28^

There remain two hurdles for mini-bulk profiling of cell-state heterogeneities. First, split-pool samples are handled somewhat primitively as negative controls—they define an aggregate null hypothesis for technical variation, which means committing many reads to a statistical cutoff.^20,26,27^ A more holistic use of these controls would be to treat them as a reference variation that compounds with the biological variation of each gene (Figures 1A and 1B).^24^ Second, when considering multiple related mini-bulk studies, comparisons are thus far limited to post hoc intersections of gene sets.^20,27,29,30^ Mini-bulk analyses should be more powerful if applied as an omnibus test with key biological variables tagged as covariates (Figure 1C). There would also be collateral benefits from such an integration. For instance, stochastic profiling^26^ is a mini-bulk approach that identifies cell-state heterogeneities using ∼25 10-cell samples and paired split-pool controls.^20^ Yet single-cell reconstruction from 10-cell mini-bulks requires 50–100 observations,^21^ a total that becomes realistic once multiple stochastic profiling studies have been properly combined.

**Figure 1.**
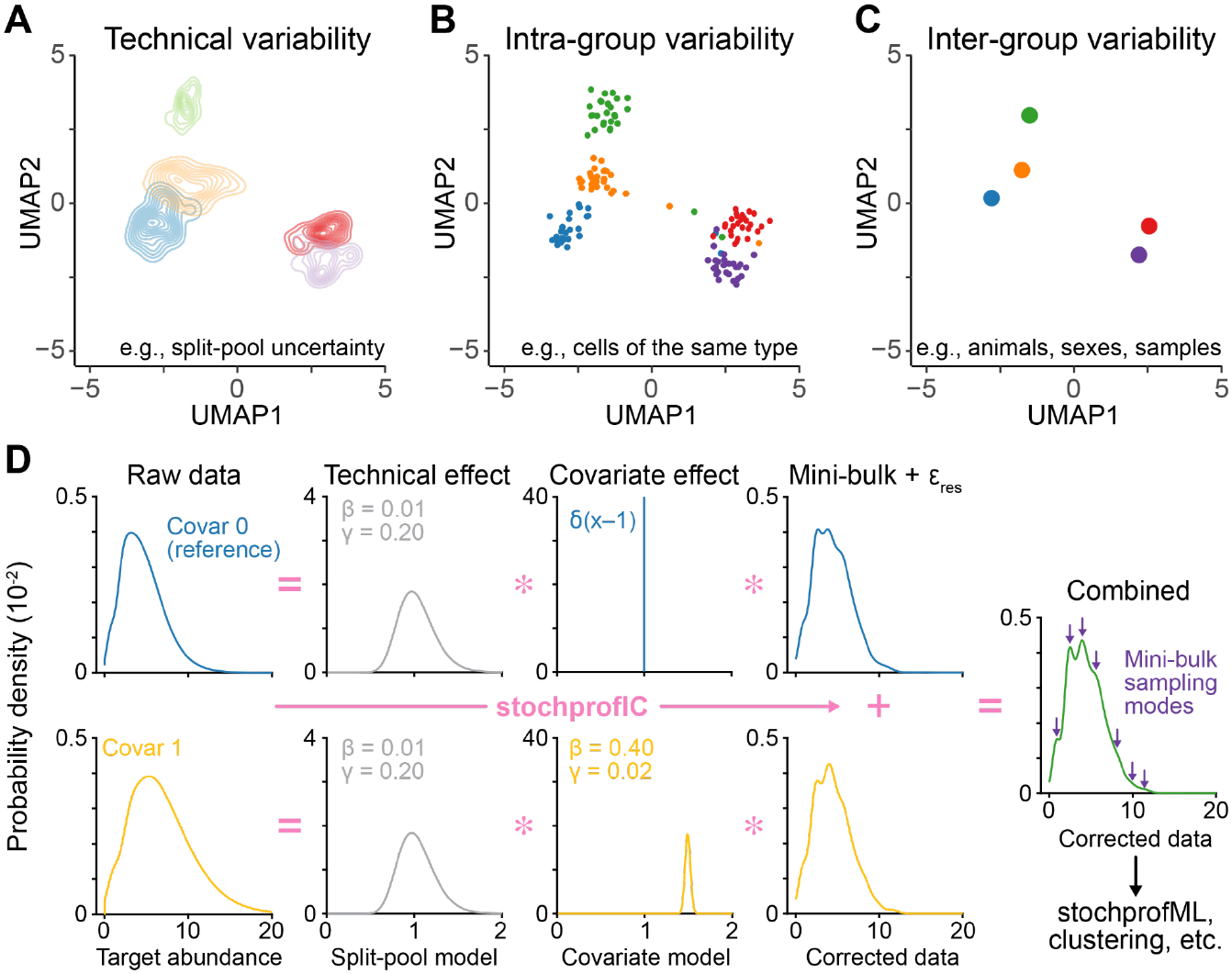
Integrating mini-bulk molecular profiles with stochprofIC. (A–C) Variation in a mini-bulk profiling study arises from (A) technical noise estimated from split-pool controls; (B) sample-to-sample fluctuations estimated from repeated mini-bulks in the same context; and (C) covariates indicating different biological contexts. Split-pool, mini-bulk, and covariate data from GSE147356 are plotted together on the same Uniform Manifold Approximation and Projection (UMAP) axes. (D) Conceptual illustration of stochprofIC. Raw data (left) are framed as a convolution of technical noise and differential covariate effects that blur mini-bulk sampling fluctuations (center). Corrected mini-bulk data from multiple biological contexts are then combined for downstream applications (right). The global log-mean and log-variance are retained here in the corrected data for clarity (in practice, they are embedded with split-pool parameters in the location and scale intercepts of the model).

In this work, we surmounted both challenges with a statistical model that formalizes technical noise and covariates as generalized additive processes, which convolve with the true biological variation of a mini-bulk sample (Figure 1D). Unlike the parsing of biological and technical (batch) variability in scRNA-seq studies,^31,32^ empirical split-pool data directly inform the model. Our two-step procedure, called stochprofIC, estimates technical noise across split-pool controls along with biological covariates across batches; then, it removes both from mini-bulk samples (Figure 1D). The combined noise- and covariate-corrected data are highly amenable to single-cell reconstruction^21^ and systems approaches more broadly.

After defining the fidelity of stochprofIC with synthetic datasets, we applied the method to identify and parameterize sex-independent state heterogeneities during early and late glioma premalignancy in mice. For multiple genes excluded previously by male–female intersection,^29^ we validated predictions of single-cell expression-state frequency by quantitative RNA FISH in independent animals. When stochprofIC was extended to a cohort of early-stage luminal breast cancers, we recovered multiple hormone receptor-associated state heterogeneities that were uniformly missed before.^20^ One transcript implicated widespread cell-to-cell differences in cargo localization when stochprofIC-corrected data were combined with a systems model of nucleocytoplasmic transport.^33^ Another that was evident in mini-bulk data but absent in scRNA-seq atlases contributed to apoptosis of luminal breast cancer organoids. As a method, stochprofIC bolsters mini-bulk studies by explicitly accounting for technical uncertainty and generalizing interpretations across similar biological settings.

## RESULTS

StochprofIC is a gene-by-gene approach that combines mini-bulk observations and matched split-pool controls. The datasets here involve 10-cell observations and controls, but the method is compatible with larger or smaller mini-bulk sizes. Stochprof<u>IC</u> uses the data to infer (I) the underlying mean–variance structure; then, it corrects (C) the observations by removing background, technical error, and estimated covariate effects from each transcript. Corrected mini-bulk observations may be directly analyzed by clustering, pathway analysis, and systems modeling or used as input for maximum-likelihood estimation of reconstructed single-cell state distributions by stochprofML.^21^

### Distributional regression of covariate and technical effects

Mini-bulk samples are typically sequenced more deeply than single cells,^14–16,34^ and some mini-bulk data types are not based on discrete counts.^13^ We thus adopted a continuous approach that posed a fully parametric, generalized additive model of location and scale (STAR Methods).^35,36^ The model uses a (log)normal probability distribution with one mini-bulk explanatory variable and an arbitrary number of covariates encoded as fixed effects. For a target (*X*) measured across *K* covariates:

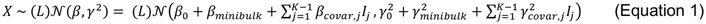

where *β* is the (log) mean and *γ*^2^ is the (log) variance, each with a split-pool-embedded intercept term (indexed as 0), a mini-bulk-associated fixed effect, and covariate-specific fixed effects (*I_j_* is an indicator function for the *j*^th^ covariate). The split-pool samples of the *K*^th^ covariate define the intercept parameters for *β* and *γ*^2^, which are shifted (*β*) or scaled (*γ*^2^) based on independent mini-bulk and covariate-specific alterations to the reference (log) mean and (log) variance. The stochprofIC package considers the transformed variance log *γ*^2^ given the link function used in the hierarchical penalized likelihood estimation algorithm.^35^ Shifts in (log) mean and scalings of (log) variance may be positive or negative, but 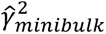 estimates should be greater than one if reflecting biological variance in excess of the technical split-pool.

After estimating terms hierarchically (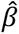 first, then 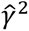) with a penalized likelihood function that assumes independence of all *β* and *γ*^2^ constituents,^35^ the target is corrected for covariates and technical effects by removing these constituents. If *X* is normally distributed:

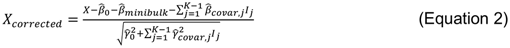

And if *X* is lognormally distributed:

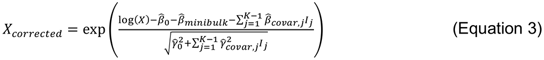

The corrected data retain the estimated mini-bulk variance along with residuals not otherwise captured by the model (Figure 1D, right).

For the correction (Equations 2 or 3) to be most valuable, mini-bulk variance must considerably exceed split-pool technical variance, which we evaluate as log *γ*^2^*_minibulk_* > 0 in the following. However, when considering *i* = 1,…, *N* targets, often the range of target abundances spans the technical noise floor of the measurement itself. We navigated the (large) *N* hypothesis tests of log *γ*^2^*_minibulk_* by using a two-step procedure (STAR Methods). First, we screened targets for robust dispersion by modeling trimmed data as a normal random variable, taking the remaining zeros as NaN, and testing the null hypothesis of log *γ*^2^*_minibulk_* = 0 at a strict familywise error rate of 5%. Then, untrimmed data from robustly dispersed targets were log-transformed and similarly analyzed without trimming and with zeros, retaining targets in which log *γ*^2^*_minibulk_* = 0 was disproved at a false-discovery rate of 15%. The procedure yielded hundreds to thousands of correctable datasets from the *N* ∼ 10^3^–10^4^ targets of a typical mini-bulk omics experiment.

### StochprofIC performance with synthetic datasets

We evaluated the suitability of stochprofIC’s additive model structure (Equation 1) with synthetic mini-bulk datasets. Starting with a representative two-state heterogeneity,^21^ we built mini-bulk samples of different sizes then convolved them with technical noise and covariate effects of different location and scale (Figure 2A and STAR Methods). Two-cell groups meet the minimum definition of a mini-bulk sample, and repeated 10-cell groups reflect frequent-to-rare cell-state heterogeneities (Figure 1D, purple), which average out in 50-cell groups.^26^ The question was whether these characteristics would be retained by stochprofIC if data were blurred by noise (including dropouts) and covariates (Figure 1D, left).

**Figure 2.**
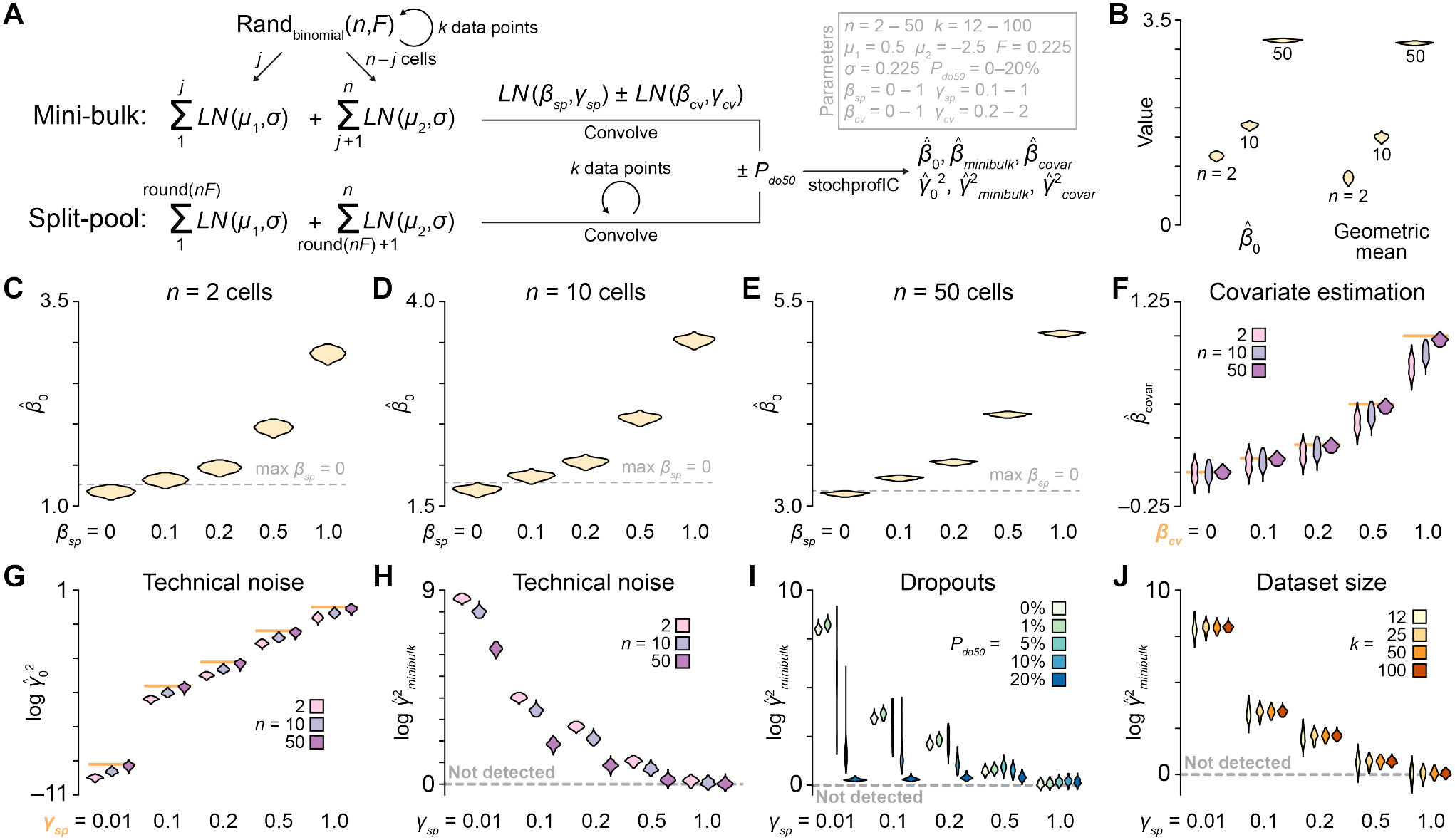
Accuracy, sensitivity, and stability assessments of stochprofIC with synthetic mini-bulk datasets. (A) Algorithm for generating synthetic data. Mini-bulk data begin with a binomial random number *j* drawn from *n* cells with a fraction (*F*) in the high expression state, which is drawn from a lognormal distribution with log mean *µ*_1_ and log standard deviation *σ*. The remaining *n* – *j* cells belong to the low expression state drawn from a lognormal distribution with log mean *µ*_2_ (< *µ*_1_) and log standard deviation *σ*. The individual draws are summed and multiplied by lognormal technical noise with log mean *β_sp_* and log standard deviation *γ_sp_*. If pertaining to the second batch, the value is additionally multiplied by lognormal covariate effects of log mean *β_cv_* and log standard deviation *γ_cv_*. The entire process is repeated *k* times. Split-pool data start with the *n*-cell average, which is convolved with technical noise and covariate effects *k* separate times to balance the overall dataset. When implemented in (I), both data types are subject to an abundance-dependent Bernoulli dropout probability before passing to stochprofIC. Fixed parameters in the gray box are taken from Bajikar et al.^21^ (B) *β^* _0_ reflects the overall geometric mean for 2-, 10-, and 50-cell mini-bulk simulations. *β_sp_* = 0, *γ_sp_* = 0.01, *β_cv_* = 0, *γ_cv_* = 0. (C–E) *β^* _0_ detects technical offsets as small as *β_sp_* = 0.1. *β_sp_* varies where indicated for *n* = 2 cells (C), 10 cells (D), and 50 cells (E). The gray dashed line marks the maximum *β^* _0_ observed when *β_sp_* = 0. *γ_sp_* = 0.01, *β_cv_* = 0, *γ_cv_* = 0. (F) *β^ _covar_* accurately captures low-to-intermediate values of *β_cv_*. *n* and *β_cv_* vary where indicated. The orange line marks the true *β_cv_* value. *β_sp_* = 0, *γ_sp_* = 0.01, *γ_cv_* = 0. (G) 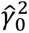 reflects *γ_sp_* in large *n* mini-bulk datasets. *n* and *γ_sp_* vary where indicated. The orange line marks the true log *γ^2^_sp_* value. *β_sp_* = 0, *β_cv_* = 0, *γ_cv_* = 0. (H) 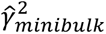 detects heterogeneous cell states for *n* ≤ 10 cells and *γ_sp_* ≤ 0.5. *n* and *γ_sp_* vary where indicated. *β_sp_* = 0, *β_cv_* = 0, *γ_cv_* = 0. (I) Dropout noise degrades, but does not eliminate, cell-state detection by 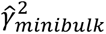. The midpoint dropout probability (*P_do50_*) and *γ_sp_* vary where indicated. *n* = 10, *β_sp_* = 0, *β_cv_* = 0, *γ_cv_* = 0. (J) 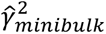 robustly detects cell states in datasets as small as *k* = 25 mini-bulk samples + 25 split-pool controls x two batches. *k* and *γ_sp_* vary where indicated. *n* = 10, *β_sp_* = 0, *β_cv_* = 0, *γ_cv_* = 0. For (H–J), the dashed line marks log 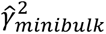 = 0, where no excess variance is estimable by stochprofIC. For (B–J), violin plots reflect the distribution of *N* = 100 independent synthetic datasets of *k* = 100 mini-bulk samples + 100 split-pool controls x two batches (B–I) or *k* = 12–100 where indicated (J). See also Figure S1.

As a reference, we began with an idealized dataset of two replicated batches each comprising 100 mini-bulk samples and 100 split-pool controls with no bias (*β_sp_* = 0) and negligible noise (*γ_sp_* = 0.01). To capture sampling uncertainty, each reference was regenerated 100 times with different binomial draws from the underlying two-state distribution (STAR Methods). The reference 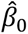 increased with mini-bulk size in proportion to the geometric mean of the reference, as expected (Figure 2B). When *β_sp_* was increased, we found that all mini-bulk groups reflected this effect as a further upward shift in 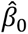 (Figures 2C–E). 50-cell groups readily discerned all *β_sp_* values, and the approximate limit of detection for 10-cell groups was *β_sp_* = 0.1, which was not reliably perceived in the two-cell groups. These changes coincided with increasingly negative 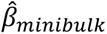 inferences, which offset the *β_sp_* bias of the split-pool samples (Figures S1A–C). With split-pool bias embedded in 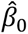 and offset as needed in 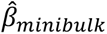, the performance towards covariate bias (*β_cv_*) was directly interpretable when applied to one of the replicated batches. The 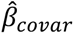 inferred by stochprofIC tracked with *β_cv_* for low-to-intermediate values tested, whereas uncertainty increased for 10- and two-cell groups, and both nominally underestimated the true value when *β_cv_* ≥ 0.5 (Figure 2F). We concluded that intermediate-to-large mini-bulks reliably capture location changes caused by technical and covariate effects.

Compared to the relationships between location and bias, we observed different trends for scale parameters in the model and noise terms in the convolution. When *γ_sp_* was increased in synthetic data, the change was readily reflected for all mini-bulk sizes as an increase in 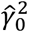, but values matched only for 50-cell mini-bulk samples (Figure 2G). The opposite was true for 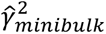, the inferred parameter most important for stochprofIC. Log *γ*^2^*_minibulk_* was detectably greater than zero for two- and 10-cell mini-bulks when *γ_sp_* ≤ 0.5, but it disappeared more rapidly for 50-cell mini-bulks due to population averaging (Figure 2H). All models poorly estimated covariate-specific noise unless it was very large (*γ_cv_* ≥ 1), when *γ_cv_* additionally had a contaminating effect on 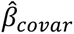 and 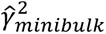 (Figures S1D–F). In practice, it is difficult to envision such a large *γ_cv_*, and thus we view it as a nuisance parameter in the model formulation.

One well recognized source of noise in scRNA-seq is technical “dropout” that arises when RNA molecules are not converted to cDNA during first-strand synthesis.^11,37^ This abundance-dependent phenomenon has analogies in mass spectrometry; therefore, we adopted an approach^38^ that models peptide dropouts as a Bernoulli random variable with a zero probability that increases sigmoidally to 100% with decreasing abundance (Figure S1G and STAR Methods). The midpoint of the sigmoid (*P_do50_*) was set at different percentiles of the 10-cell synthetic datasets, and dropped-out data were rerun through stochprofIC with a focus on 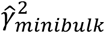. When *γ_sp_* was low, even very small values of *P_do50_* degraded the magnitude of 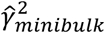 (Figure 2I). (This sensitivity could be addressed by using a zero-adjusted lognormal^36^ instead of Equation 1, but it would add a third distributional parameter to the estimation problem.) Interestingly, when *γ_sp_* was intermediate and around the likely precision of small-sample transcriptomic methods (*γ_sp_* = 0.5), we found that 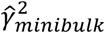 was much more resilient, remaining detectable up to *P_do50_* = 20%. On a per-cell basis, transcripts should be less likely to drop out with mini-bulking; nonetheless, the results here indicate that the Equation 1 formulation of stochprofIC has considerable robustness to dropout noise.

As a final assessment, we asked how many mini-bulk observations (*k*)—balanced across samples, split-pools, and covariates (Figure 2A)—were needed for stochprofIC to retain its performance with respect to 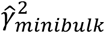. Taking *γ_sp_* = 0.5 as a likely estimate of technical precision, we found that stochprofIC reliably discriminates cell states with at least 25 mini-bulk samples plus 25 split-pool controls collected in two batches. These numbers are highly concordant with the designs of mini-bulk approaches like stochastic profiling^26^ and stochprofML^21^ (*k* ≥ 25 in two batches). Our overall interpretation is that stochprofIC reliably extracts cell-state variations as 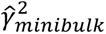 from intermediate-to-small mini-bulks when studies of this size are combined and covariate-specific noise is not exaggerated.

### Application to early and late glioma premalignancy

To compare stochprofIC against competing alternatives, we sought existing mini-bulk datasets with split-pool controls and well-defined covariates. In an inducible murine model of gliomagenesis,^39^ Sutcliffe et al.^29^ performed repeated mini-bulk profiling of tdTomato (tdT)-positive mutant oligodendrocyte precursor cells (OPCs; Figure 3A). The study considered two separate contexts: i) ∼1 week after a five-day induction (“12 dpi”), when cells have responded to the abrupt loss of *Nf1* and *Trp53* tumor suppressors; and ii) ∼3 months after induction (“90 dpi”), when mutant cells have fractionally (de)differentiated along various lineages. Each context was mini-bulk profiled in males and females (GSE147360), recognizing the sexual dimorphism of advanced gliomas.^40^ Our goal here was to correct for sex as a covariate and split-pool technical noise in the two pre-glioma contexts by using stochprofIC.

**Figure 3.**
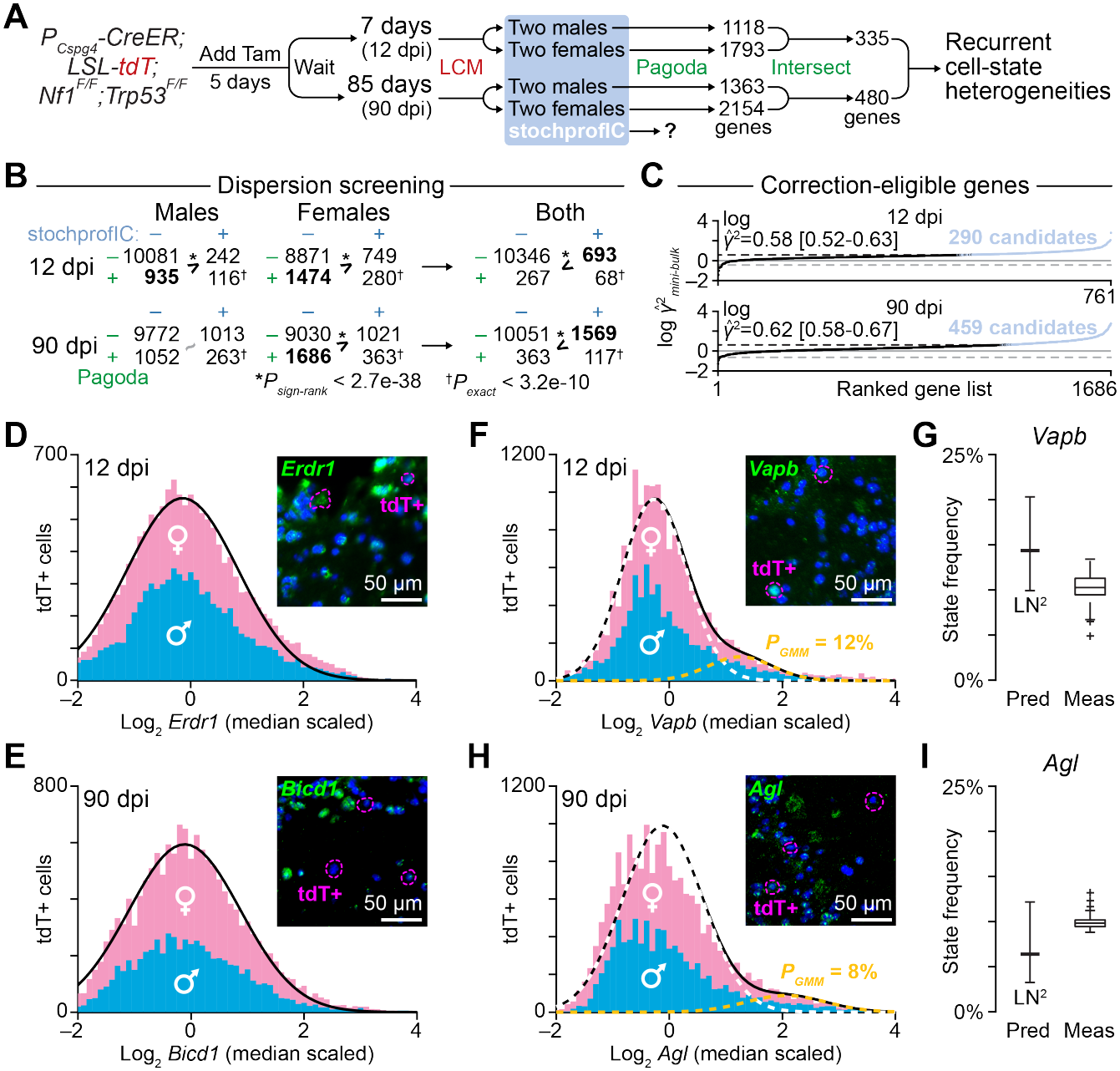
StochprofIC enhances identification and quantitation of sex-independent expression-state heterogeneities during murine gliomagenesis. (A) Sutcliffe et al.^29^ used mice engineered with estrogen-regulated Cre under the *Cspg4* promoter, loxP-STOP-loxP tdTomato (tdT), and floxed alleles of *Nf1* and *Trp53* (*P_Cspg4_-CreER; LSL-tdT; Nf1^F/F;^ Trp53^F/F^*). Cre was induced in *Cspg4*-expressing oligodendrocyte precursor cells (OPCs) with five daily doses of tamoxifen (Tam), and then tdTomato-positive OPCs were isolated at the indicated times by laser capture microdissection (LCM; *N* = two males + two females at each time point). The study identified genes in males or females that were overdispersed given their abundance using Pagoda^41^ and then intersected these genes by sex. (B) Comparison of dispersion calls between Pagoda (rows) and stochprofIC (columns) for each sex or both sexes at the indicated time points. Enrichments between the two methods were assessed by hypergeometric test and off-diagonal biases by sign-rank test. (C) Genes with robust dispersion were ranked by 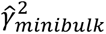, and genes with log *γ*^2^*_minibulk_* > 0 (*q* < 0.15) were identified as candidates (blue). A log 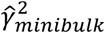 threshold for each timepoint was estimated by smoothing candidate calls with a 100-gene moving window. The midpoint 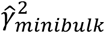 threshold is shown with 5th and 95th percentiles in brackets. (D and E) *Erdr1* at 12 dpi (D) and *Bicd1* at 90 dpi (E) have one-state log-normal abundance in tdT+ OPCs from males (blue) and females (pink). (F–I) *Vapb* at 12 dpi [(F) and (G)] and *Agl* at 90 dpi [(H) and (I)] have two-state log-normal abundance in tdT+ OPCs from males (blue) and females (pink). In (G) and (I), the measured (Meas) frequency of cells in the high expression state is compared to the predicted (Pred) frequency from stochprofML^21^ given stochprofIC-corrected data and a two-state log-normal model with equal variance (LN^2^). For (D–F) and (H), data are from *N* = 13977 (D), 4879 (E), 16229 (F), and 6278 (H) tdT+ OPCs in four mice (two males, two females) at the indicated time points. The inset shows representative RNA FISH intensities (green) in the olfactory bulb tissue with tdT+ OPCs outlined in magenta. The scale bar is 50 µm. For (G) and (I), predictions are shown as the maximum-likelihood estimate ± 95% CI and measurements are shown as the estimated frequency from *N* = 50 bootstrap replications of the starting total-cell population for each gene. See also Figures S2 and S3 and File S1.

The prior study^29^ was similarly interested in identifying cell-state heterogeneities that recurred between sexes. This was achieved by identifying biologically overdisperse transcripts and intersecting the results from males and females (Figure 3A). Gene overdispersion was quantified with Pagoda,^41^ which builds a transcriptome-wide regression model of observed *µ*^2^/*γ*^2^ versus *µ* to calculate an “adjusted variance” relative to expectation for a given abundance. In Sutcliffe et al.,^29^ split-pool controls created a null distribution for transcriptome-wide adjusted variance, thereby defining a cutoff for adjusted variance in the corresponding mini-bulk samples. We applied this approach to the complete sex-separated datasets (STAR Methods) and identified a few thousand overdispersed transcripts each in males and females, along with several hundred that recur in both (Figure 3A and File S1).

As implemented, Pagoda is most akin to the front-end dispersion screen of stochprofIC. The key difference is that Pagoda builds a transcriptome-wide null distribution of dispersion from split-pool samples, whereas stochprofIC makes gene-specific comparisons. Conceptually, Pagoda should have a more reliable null reference with smaller sample sizes (e.g., one sex), because the entire measured transcriptome contributes to the null distribution. With more split-pool samples, however, the Pagoda reference is unlikely to change much, whereas more observations will solidify the *β^* _0_ and 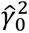 estimates of stochprofIC. We asked whether this inflection in sensitivity was evident when combining the 28 mini-bulk samples and 20 split-pool controls each from males and female OPCs in GSE147360.

We ensured a fair comparison across methods, subsets, and supersets by considering only the transcripts that passed standard Pagoda filtering criteria when both sexes were analyzed individually (11,374 genes at 12 dpi and 12,100 genes at 90 dpi; STAR Methods). Dispersion screening was performed on transcripts per million (TPM) estimates with stochprofIC, and the genes with *γ*^2^*_minibulk_* > 0 were compared with the adjusted variance-based dispersion calls of Pagoda. Consistently, there was significant overlap between the two methods (*P* < 1e-20 by Fisher’s exact test), but Pagoda identified many more overdispersed genes in three of four cases (*P* < 3e-38 by binomial test; Figure 3B, left). We then combined the sex data by intersection (Pagoda) or modeling of *β_sex_* and *γ*^2^*_sex_* (stochprofIC) and found that the bias flipped, with 2.3- and 3.5-fold more genes retained by stochprofIC at 12 dpi and 90 dpi respectively (Figure 3B, right). Therefore, at the scale of this study, sensitivity improvements emerge when stochprofIC considers as few as one binary covariate.

For the genes with robust dispersion, we asked whether excess mini-bulk variance was detectable after log transformation of all observations (56 mini-bulk samples + 40 split-pool controls) and including sex as a covariate. At both timepoints, log 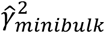 skewed strongly positive, and false-discovery cutoffs were surpassed when the excess mini-bulk log variance was greater than ∼0.6 (Figure 3C). 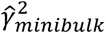 estimates were anti-correlated with 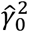 (Figures S3A and S3B), reflecting expectation that precisely measured genes would have more detectable biological variation. However, 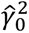 did not act as a strict cutoff, because some technically precise genes did not have detectable *γ*^2^*_minibulk_* (e.g., *Tmem170*; Figure S3A), whereas some imprecisely measured genes did (e.g., *Rgs4*; Figure S3A). We note that even upon mini-bulking, the percentage of OPC genes with variance above measured technical noise is small. A similar situation likely exists for scRNA-seq yet technical noise of this kind is rarely measured.

After correction of split-pool- and sex-specific effects, we passed the remaining 12-dpi and 90-dpi candidates to stochprofML.^21^ This package uses mini-bulk data to test three single-cell models: i) a one-state log-normal distribution (LN) as the null hypothesis versus different two-state mixtures of log-normal distributions with ii) equal variance (LN^2^) or iii) unequal variance (rLN^2^; STAR Methods). We ranked models by Bayesian Information Criterion weights^42^ (BICw; File S1) and selected five genes per time point for further study, prioritizing different classifications by Pagoda and stochprofIC (Figure 3B). Predicted distributions of transcript abundance were tested by RNA fluorescence in situ hybridization (FISH) combined with tdT immunofluorescence for OPCs and digital image segmentation (Figure S2; STAR Methods). We assessed generality at the 12 and 90 dpi time points by analyzing two males and two females each that were not mini-bulk profiled in the original study.^29^

At 12 dpi, one-state LN models were heavily favored for *Erdr1* (BICw = 0.98) and *Echs1* (BICw = 0.94) after stochprofIC correction of mini-bulk data; neither gene was overdisperse in males or females by Pagoda. Accordingly, per-OPC RNA FISH intensities for *Erdr1* and *Echs1* were well fit by a single log-normal distribution (Figures 3D and S3C). At 90 dpi, we observed the same for *Bicd1* (LN BICw = 0.81; Figure 3E) and *Eml1* (Figure S3D). The latter gene was notable because LN was only marginally preferred in stochprofML (BICw = 0.51), and Pagoda identified *Eml1* as overdisperse in both sexes (File S1). Although *Eml1* is recurrently variable at 12 dpi, the stochprofIC-corrected data and RNA FISH support a single mode of cell-state regulation.

We applied the same methods to three 12-dpi genes and three 90-dpi genes where two-state models were preferred, substituting a two-population Gaussian mixture model to fit the RNA FISH data (STAR Methods). For the 12-dpi genes *Vapb* (LN^2^ BICw = 0.75) and *C1qtnf4* (LN^2^ BICw = 0.79), stochprofML predicted high-state expression frequencies of 14% (95% CI: 9.9–20%) and 9.8% (95% CI: 6.7–14%), respectively, which agreed well with the bootstrapped medians of 10% and 7.5% by RNA FISH (Figures 3F, 3G, S3E, and S3F; STAR Methods). Predictions likewise coincided for the 90-dpi genes *Agl* (LN^2^ BICw = 0.64) and *Myo9a* (LN^2^ BICw = 0.84), with inferred frequencies of 6.4% (95% CI: 3.2–12%) and 9.4% (95% CI: 6.5–13%) that overlapped the bootstrapped measured frequencies of 9.8% and 10% (Figures 3H, 3I, S3G, and S3H). Some genes were more nuanced in OPCs. Corrected mini-bulk measurements of the 12-dpi gene *Hdac4* (LN^2^ BICw = 0.74), for example, led to a stochprofML-predicted expression frequency of 11% (95% CI: 7.7–17%; Figures S3I and S3J). Although the median RNA FISH frequency for *Hdac4* was 32%, we noted multiple bootstrap replications between 13–17% consistent with the prediction (Figure S3J, yellow) that suggested competing local minima in the two-state Gaussian fit. The 90-dpi gene *Inpp1* was challenging, because stochprofML could not distinguish whether its two-state variance should be equal (LN^2^ BICw = 0.47) or unequal (rLN^2^ BICw = 0.51). Nevertheless, the inferred *Inpp1* expression frequency was comparable: 10% (95% CI: 7.2–14%) for LN^2^ and 12% (95% CI: 8.0–18%) for rLN^2^. We split the tie with Occam’s razor and fit the RNA FISH observations to LN^2^, finding a somewhat elevated bootstrapped median frequency of 15% (Figure S3K and S3L). The slight difference is attributable to the greater tendency for male OPCs to occupy the *Inpp1*-high state (Figure S3K, yellow); stochprofML does not handle covariates like stochprofIC. Taken together across ten OPC genes at two very different time points of gliomagenesis, we conclude that stochprofIC harmonizes covariate-dependent mini-bulk data, enabling accurate single-gene predictions of state heterogeneity by stochprofML.

### Application to early-stage luminal breast cancer

We next applied stochprofIC to a mini-bulk study where covariates (patients) were more numerous and the effect size between them more pronounced. Singh et al.^20^ mini-bulk profiled the carcinoma cells of five late-onset, early-stage luminal breast cancers (GSE147356). Two of these cases were so infiltrated with stroma cells that KRT8-guided microdissection was required for isolation (UVABC3 and UVABC5; Figure 4A). Heterogeneity within the cohort was further reflected in the results of the Pagoda-based dispersion analysis: among the 5982 genes identified in any single tumor, only seven were identified in all five. Singh et al.^20^ thus adopted a majority-vote approach, defining a cell-state heterogeneity as recurrent if it appeared in at least 50% of cases (Figure 4A). We asked whether stochprofIC (plus stochprofML) would provide an enhanced perspective on recurrent cell states after correcting for technical noise and patient as a covariate.

**Figure 4.**
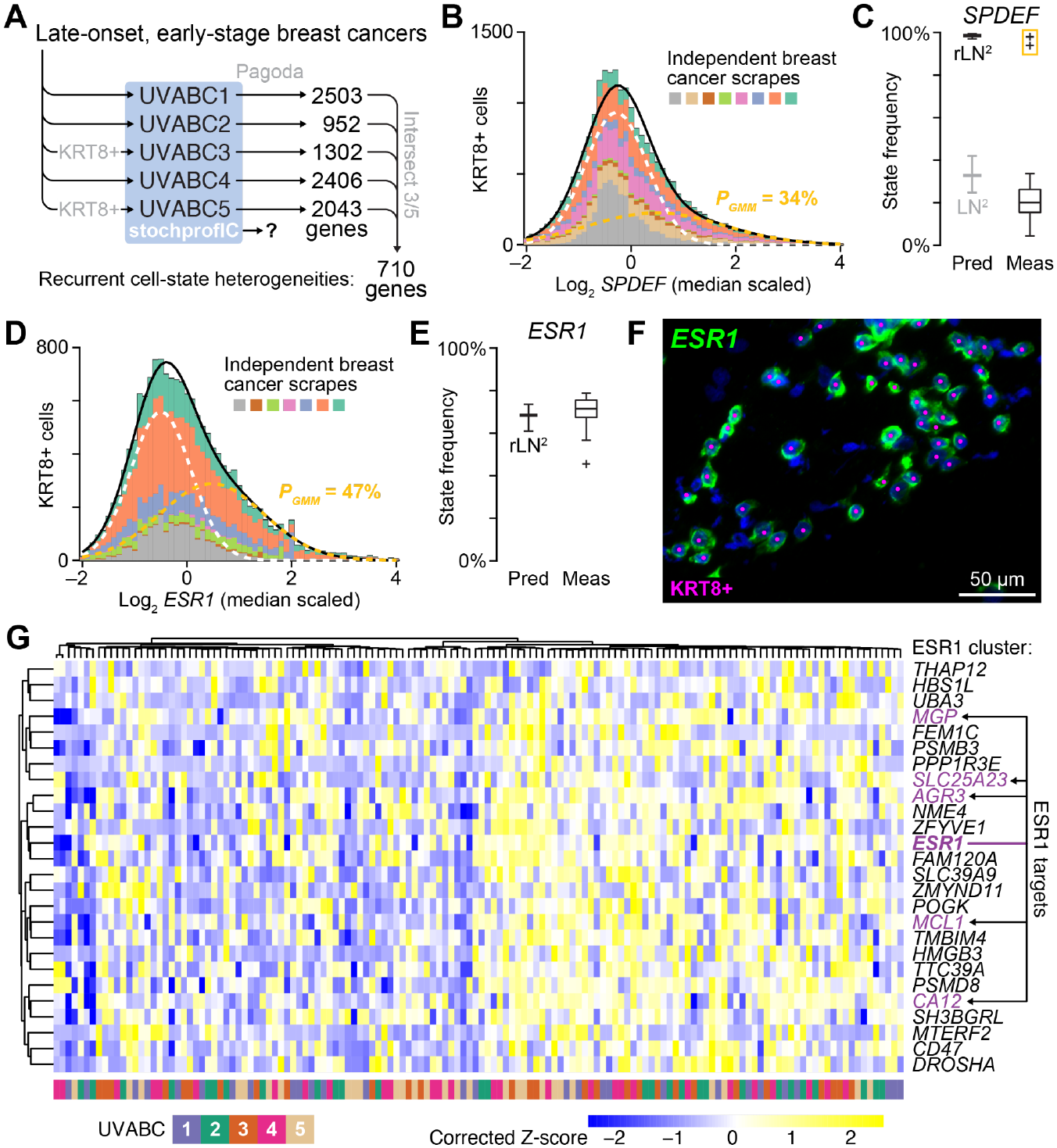
StochprofIC recovers cell states for hormone-dependent transcription factors and target genes in luminal breast cancer. (A) Singh et al.^20^ profiled five late-onset, early-stage breast cancers by mini-bulk LCM that was guided, when needed, by KRT8 immunofluorescence (UVABC3 and UVABC5). The study identified genes in single patients that were overdispersed given their abundance using Pagoda^41^ and then intersected these genes to find candidates that recurred in most (= ⅗) cases. (B and C) *SPDEF* has a two-state log-normal distribution with unequal variance in KRT8+ breast cancer cells from *N* = 8 independent cases (colors; File S2). In (C), the measured (Meas) frequency of cells in the high expression state is compared to the predicted (Pred) frequency from stochprofML^21^ given stochprofIC-corrected data and two-state log-normal models with equal variance (LN^2^, gray) or unequal variance (rLN^2^, black). Two measured bootstrapped replications with high expression frequency are boxed in yellow. (D–F) *ESR1* has a two-state log-normal distribution with unequal variance in KRT8+ breast cancer cells from *N* = 7 independent cases (colors; File S2). In (E), the measured (Meas) frequency of cells in the high expression state is compared to the predicted (Pred) frequency from stochprofML^21^ given stochprofIC-corrected data and a two-state log-normal model with unequal variance (rLN^2^). The image in (F) shows representative RNA FISH intensities for *ESR1* (green) in a tumor scrape (UVABC214; File S2) with KRT8+ breast cancer cells dotted in magenta. The scale bar is 50 µm. (G) Extracted row-standardized clustergram of stochprofIC-corrected data (Figure S4B) containing *ESR1* and multiple known ESR1 target genes (purple). Integration of UVABC mini-bulk samples is illustrated by the column-clustered patient track at the bottom. For (C) and (E), predictions are shown as the maximum-likelihood estimate ± 95% CI and measurements are shown as the estimated frequency from *N* = 50 bootstrap replications of the starting total-cell population for each gene. See also Figures S2 and S4 and File S2.

Upon integrating 5 x 28 = 140 mini-bulk samples and 5 x 20 = 100 split-pool controls with stochprofIC, we identified 488 genes with detectable *γ*^2^*_minibulk_*. Clustering of this subset before and after correction illustrated the patient-specific mini-bulk transcriptomes and their resolution with stochprofIC (Figures S4A and S4B). We downsampled the data and observed linear decreases in the number of candidates with fewer mini-bulk samples and fewer samples plus controls (Figure S4C). Unlike the majority-vote approach,^20^ there was no evidence of saturation when using the complete dataset from five cases, suggesting *γ_sp_* ≥ 0.5 for many genes (Figure 2J). The candidate genes were tested with stochprofML, and 86% (= 421 / 488) conformed to a two-state distribution (File S2). Remarkably, 62% (= 260 / 421) of the genes were not detected by Pagoda as overdisperse for any single UVABC case in the prior analysis.^20^ These new recurrent expression states were pursued for validation and follow up; here, we focused less on markers and more on pathways and mechanisms.

One prominent gene on the list was the transcription factor *SPDEF* (File S2), an important effector of luminal differentiation in breast tissue and in breast cancers.^43^ We found that stochprofML strongly excluded a one-state distribution for *SPDEF* (LN BICw = 8.7e-6). However, the evidence was somewhat split between rLN2 (BICw = 0.73), with a predicted expression frequency of 99%, and LN2 (BICw = 0.27), with a predicted expression frequency of 33%. To clarify, we obtained tumor scrapes from an independent set of luminal breast cancers (File S2), hybridized for *SPDEF*, and co-identified carcinoma cells by KRT8 immunostaining (STAR Methods). Using the same gating and quantification as for tdT+ OPCs, we recurrently detected a right-skewed distribution of *SPDEF* in KRT8+ breast cancer cells (Figures 4B and S4D). Interestingly, the bootstrapped median frequencies of *SPDEF* by RNA FISH straddled the divergent frequencies predicted by rLN2 and LN2 (Figure 4C), suggesting two best fits that were hardly discriminable. *SPDEF* abundance is modulated by multiple upstream transcription factors including FOXA1, GATA3, and ESR1,^43^ suggesting why it might not neatly form a two-state distribution. Notwithstanding, the concordant bimodality of prediction and measurement supported that the underlying generating function for SPDEF was distilled by stochprofIC.

The luminal-defining transcription factor *ESR1* was also identified as a recurrent two-state heterogeneity in the new analysis, with an rLN^2^ (BICw = 0.98) expression frequency of 68% (95% CI: 61–74%). Using scrapes from the same luminal breast cancer as before, we quantified *ESR1* in KRT8+ cells and estimated a median bootstrapped expression frequency of 72% (Figures 4D–F), corroborating the prediction. Previously, Singh et al.^20^ detected ESR1 action indirectly as an enrichment of its target genes in the Pagoda-based list (Figure 4A); stochprofIC created an opportunity to inspect its correlates directly. We extracted the stochprofIC-corrected hierarchical cluster surrounding *ESR1* and readily identified five recognized targets: *MGP*,^44^ *SLC25A23*,^45^ *AGR3*,^46^ *MCL1*,^47^ and *CA12*^48^ (Figures 4G and S4B). Of these, only *AGR3* and *CA12* were among the 710 genes identified as recurrent by Pagoda (Figure 4A). These results show how stochprofIC increases the statistical power and biological interpretability of mini-bulk studies by reconciling disparate-but-related samples (Figures S4A and S4B).

### Systems modeling with stochprofIC outputs

A surprising gene in the luminal breast cancer list was the GTPase *RAN*. Normally thought of as a housekeeping gene, *RAN* was instead favored as rLN^2^ (BICw = 0.53) with an expression frequency of 62% (95% CI: 50–68%). This nearly matched the 61% median bootstrapped expression frequency of *RAN* measured in KRT8+ cells of tumor scrapes (Figures 5A–C). Moreover, we noted a pronounced bimodality in some scrapes (Figure 5A, gray and tan), suggesting functional differences given the critical role for RAN in nucleocytoplasmic transport.^49^ To address this question computationally, we leveraged an existing model (MODEL2210060001) of nucleocytoplasmic transport involving RAN and ten other key species. Starting with its prior calibration for a breast epithelial line,^33^ we perturbed initial conditions according to the uncorrected or stochprofIC-corrected transcript abundances of the relevant genes (Figure 4D; STAR Methods). (Note that stochprofIC corrections are possible even when log *γ*^2^*_minibulk_* is not significantly greater than zero, as was the case for all species in the model other than RAN.) The initialized systems were run to steady state and then challenged with a bolus of cargo containing either i) a nuclear localization sequence (NLS), ii) a tighter bivalent NLS (2xNLS), or iii) a direct importin-β-binding (IBB) sequence, each with or without a nuclear export sequence (NES). The 3 x 2 = six models were again run to steady state and the nuclear-to-cytoplasmic (N/C) ratio was calculated for the 280 (= 2 x 140) initializations in Figure 4D to test for heterogeneities in N/C shuttling state.

**Figure 5.**
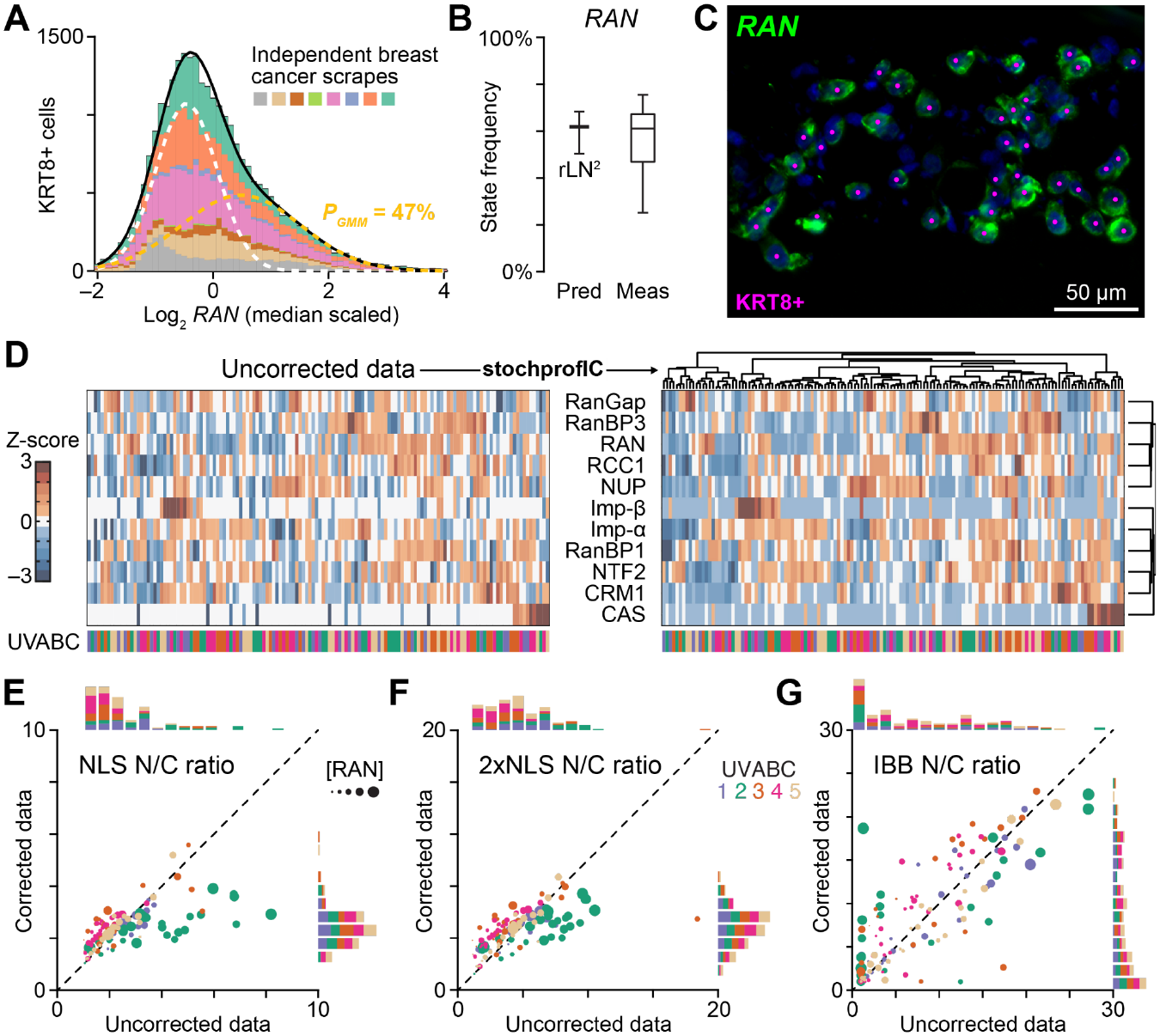
StochprofIC-informed systems modeling of nucleocytoplasmic transport. (A–C) *RAN* has a two-state log-normal distribution with unequal variance in KRT8+ breast cancer cells from *N* = 8 independent cases (colors; File S2). In (B), the measured (Meas) frequency of cells in the high expression state is compared to the predicted (Pred) frequency from stochprofML^21^ given stochprofIC-corrected data and a two-state log-normal model with unequal variance (rLN^2^). Predictions are shown as the maximum-likelihood estimate ± 95% CI and measurements are shown as the estimated frequency from *N* = 50 bootstrap replications of the starting total-cell population. The image in (C) shows representative RNA FISH intensities for *RAN* (green) in a tumor scrape (UVABC209; File S2) with KRT8+ breast cancer cells dotted in magenta. The scale bar is 50 µm. (D) Row-standardized clustering of log_2_-transformed, transcript-aggregated species used to scale initial conditions for a systems model of nucleocytoplasmic shuttling (MODEL2210060001). Data are shown for the original uncorrected data^20^ (zeros substituted with the row median; left) and after stochprofIC correction (right). UVABC mini-bulk samples are illustrated by the column-clustered patient track at the bottom. (E–G) Predicted steady-state nuclear-to-cytoplasmic (N/C) ratio of the indicated nuclear-localized cargo when uncorrected initial conditions (x-axis) or stochprofIC-corrected initial conditions (y-axis) are used in the model. UVABC cases are color-coded as in (D), and marker size is proportional to the square root of uncorrected RAN abundance. See Wang et al.^33^ for further details on cargo and the model. See also Figures S2 and S5 and File S2.

We correlated the N/C predictions with uncorrected and corrected initial conditions for each cargo and made several observations. With uncorrected initial conditions, UVABC2 was a clear outlier for NLS-containing cargo, regardless of whether the cargo additionally contained an NES (Figures 5E, 5F, S5A, and S5B). The effect was less pronounced for 2xNLS, but in either setting the uncorrected distributions suggested a right-tailed heterogeneity (top histograms). By contrast, the distributions of NLS-containing cargo with corrected initial conditions were more well mixed and Gaussian (right histograms), which suggested nominal N/C state differences after accounting for covariates.

A very different N/C picture emerged for IBB cargo (Figures 5G and S5C). Individual models were more dispersed overall, and UVABC2 revealed itself as an outlier differently, with a skew toward lower N/C ratios (top histogram). Importantly, although stochprofIC correction improved the mixing of N/C states across UVABC cases, the strong rightward skew persisted (right histogram), suggesting different shuttling characteristics of IBB-like cargo in luminal breast cancers. Whereas stochprofIC identifies expression-state heterogeneities of single genes when combined with stochprofML, it proposes emergent cell-state heterogeneities when combined with systems models.

### Phenotypic validation of a rare MAP3K11-high state identified by stochprofIC

Last, we asked whether some candidate transcripts define cell states relevant to breast tumorigenesis by examining *MAP3K11*, a mitogen-activated protein kinase kinase kinase nominated exclusively by stochprofIC. *MAP3K11* garnered attention because two related genes, *MAP3K1* and *MAP2K4*, are recurrently mutated in luminal breast cancer.^50^ Like MAP3K1, MAP3K11 is activated by inflammatory ligands and transduces stress signals through MAP2K4,^51^ suggesting it may be a tumor suppressor inactivated by mechanisms other than mutation.^20,52,53^ *MAP3K11* is also representative of the “dim transcriptome” returned by scRNA-seq—it is detected in only 16% (= 16221 / 100064) and 5.4% (= 89513 / 1666717) of cells from breast cancer^6^ and normal breast tissue^54^ atlases, respectively. Moreover, only 0.13% (= 135 / 100064) and 0.0029% (= 48 / 1666717) of cells have more than five counts. This contrasts *MAP3K11*’s bulk expression in breast tissue (63 median transcripts per million [TPM; IQR: 52–72 TPM]; *N* = 514 samples from GTEx^55^) and mini-bulk expression in breast cancer (15 median TPM [IQR: 6.5–36 TPM]; *N* = 240 samples from Singh et al.^20^).

The stochprofML–RNA FISH validation of *MAP3K11* returned a rLN2 (BICw = 0.64)-predicted expression frequency of 9.0% (95% CI: 6.3–16%), which nearly matched the measured bootstrapped value of 10% (Figure 6A–C). MAP3K11 expression and heterogeneity was also evident at the protein level in a separate cohort^56^ of normal luminal breast epithelia and ESR1+ luminal breast cancers (Figure S6A and STAR Methods). With the *MAP3K11*-high state verified, we considered its implications for luminal breast cancer. There is evidence that the ESR1 ligand β-estradiol inhibits MAP3K11 posttranslationally,^57^ and *ESR1* itself is heterogeneous in most luminal carcinomas (Figure 4D–F). Yet, in the stochprofIC-corrected data of Singh et al.,^20^ *MAP3K11* was uncorrelated with *ESR1* (log Pearson *R* = 0.01, *P* = 0.94). We hypothesized that the resulting spectrum of MAP3K11 activity states might relate to stress-induced cell death based on past work.^52,57^

**Figure 6.**
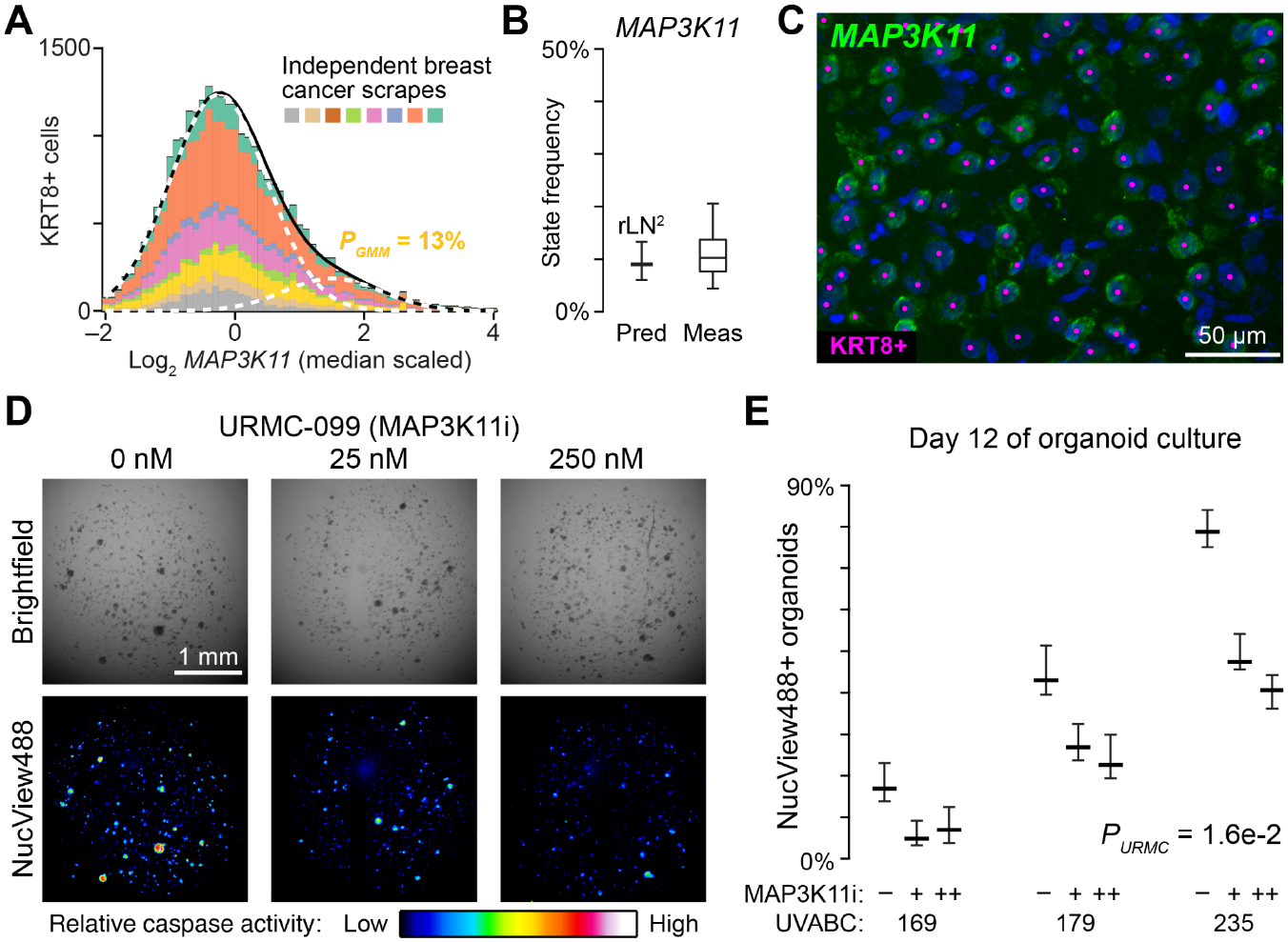
A rare *MAP3K11*-high state identified by stochprofIC regulates apoptosis in luminal breast cancer cells. (A–C) *MAP3K11* has a two-state log-normal distribution with unequal variance in KRT8+ breast cancer cells from *N* = 8 independent cases (colors; File S2). In (B), the measured (Meas) frequency of cells in the high expression state is compared to the predicted (Pred) frequency from stochprofML^21^ given stochprofIC-corrected data and a two-state log-normal model with unequal variance (rLN^2^). Predictions are shown as the maximum-likelihood estimate ± 95% CI and measurements are shown as the estimated frequency from *N* = 50 bootstrap replications of the starting total-cell population. The image in (C) shows representative RNA FISH intensities for *MAP3K11* (green) in a tumor scrape (UVABC218; File S2) with KRT8+ breast cancer cells dotted in magenta. The scale bar is 50 µm. (D and E) MAP3K11 inhibition with URMC-099 decreases spontaneous apoptosis of zero-passage luminal breast cancer organoids. (D) Brightfield image (upper) of an organoid culture (UVABC235; File S2) treated with the indicated concentration of URMC-099 on day 4 and imaged with NucView488 live-cell caspase reporter (lower) on day 12. The scale bar is 1 mm. (E) Quantification of NucView488+ organoid frequencies from *N* = 3 independent patients. Data are shown as the mean ± 84% bootstrapped confidence intervals,^70^ and the fixed effect of URMC-099 (*P_URMC_*) was assessed as a continuous factor by multiway ANOVA of arcsine square-root transformed percentages with UVABC case as an additional fixed effect. See also Figures S2 and S6 and Files S2 and S4.

To test the phenotypic impact of ablating a MAP3K11-high state, we used “zero-passage” organoids derived directly from luminal breast cancer scrapes.^58^ The initial culture as organoids is very stressful for most samples, with cells dying to variable degrees as others proliferate. We reasoned that zero-passage cultures would be a good setting to evaluate the response, if any, to acute MAP3K11 inhibition with the selective inhibitor, URMC-099.^59^ Luminal breast cancer cells were isolated from patients and cultured as organoids, with URMC-099 added at different concentrations on day 4. For three independent cases (File S2), we observed consistent paired decreases in organoid apoptosis with both doses of URMC-099 from day 8 and and past day 12 (Figures 6D, 6E, and S6B). In one case treated with high-dose URMC-099 (UVABC179), apoptosis decreased even though overall cell death increased, supporting that the apoptotic phenotype is specific (Figures 6E and S6C). For MAP3K11, perturbing a relatively rare cell state (Figures 6A–C) is sufficient to elicit a vintage hallmark of cancer.^60^

## DISCUSSION

StochprofIC formalizes how explanatory variables of a mini-bulk study contribute to the distributional properties of measured gene profiles. If split-pool controls are well paired with mini-bulk samples, then inferred shifts in log mean should mostly reflect fixed covariate effects between study groups. Reciprocally, scaling of log variance ought to be largely through *γ*^2^*_minibulk_*, with *γ*^2^*_covar_* capturing residual batch effects of day-to-day processing (Equation 1). This compartmentalization of location and scale makes controlled mini-bulk studies more informative both technically and biologically.

The estimates of technical and excess biological variance are worth revisiting. In studies that combine 100 (Sutcliffe et al.^29^) to 250 (Singh et al.^20^) repeated observations of a cell type, we detect only a few hundred genes with *γ*^2^*_minibulk_* > 0 when the 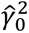 of split-pool controls is considered. Such empirical estimates of technical noise have disappeared with the commercialization of scRNA-seq. Older plate-based scRNA-seq algorithms like Pagoda^41^ conveyed measurement uncertainty by modeling it as inversely proportional to the measured abundance. Now, foundation models for droplet-based scRNA-seq ask AI to learn an analogous relationship from different tokenizations^61,62^ of individual cells. None use direct measurements of technical uncertainty, because the data are largely absent in the field (as noted in Fang et al.^25^). One path ahead for scRNA-seq may be to build “pseudo-mini-bulks” of ∼10 cells within a type and compare with “pseudo-split-pools” of ∼1000 cells where cell-state variations are averaged away.^16,26^ These synthetic mini-bulk studies would be immediately compatible with stochprofIC, much like the fully synthetic data used here to assess model performance.

The noise and covariate corrections of stochprofIC have implications for applying systems biology models at scale.^63^ Bulk transcriptomics in human samples have initialized patient-specific models,^64,65^ and scRNA-seq-based models are piloting for different applications.^66,67^ To inform these models, corrected mini-bulk data from stochprofIC provide an alternative source of generalized biological variability. If single-cell data are essential, specific genes may be sampled from the distributions inferred by stochprofML and combined using gene-to-gene covariance estimates from the corrected mini-bulk profiles. Systems biologists agree that models of biological mechanisms are best constrained by measurements that recognize the data-generating mechanisms.^68^

## Limitations of the study

As a package, stochprofIC automatically handles covariates as categorical fixed effects. Modeling random effects, mixed effects, and nested effects is possible, but it requires modifying the source code for a specific covariate structure of interest. Throughout this work, we validate predictions of expression frequency by overlapping uncertainty intervals of a stochprofML model with those of RNA FISH data, but “not significantly different” does not mean a perfect match. What gives confidence is the overall concordance correlation coefficient (= 0.987) between predictions and measurements for 10 genes in three biological contexts (Figures 3F–I, 4B–E, 5A–C, 6A–C, and S3E–L). Finally, a cell-state heterogeneity may be recurrent, accurately predicted, and experimentally validated, but that does not mean the two states are biologically important. Genetic manipulations^69^ that homogenize an endogenous two-state heterogeneity are ultimately required to assert functional relevance. The pharmacologic perturbation of MAP3K11 with URMC-099 is a first step in this direction.

## ACKNOWLEDGMENTS

We thank Jungeun Kim for assistance with mouse samples, Piotr Przanowski for early testing of stochprofIC code, and Aakrosh Ratan for reviewing the manuscript. This work was supported by grants from the National Institutes of Health (R01-CA256199 to K.A.J. and H.Z., U01-CA215794 to K.A.J. and C.F. [approved foreign component], R01-CA194470 to K.A.J. and H.Z., U54-CA274499 to K.A.J., H.Z., and S.L.S.; R50-CA265089 to L.W.; and K00-CA253732 to R.K.P.) and the Coulter Foundation (AC10854 to K.A.J. and S.L.S.), as well as fellowship support from the National Institutes of Health (T32-GM145443 to K.A.J. and R.B.H.; T32-GM156694 to R.B.H. and N.L.; T32-GM136615 to S.A.V.) and the National Science Foundation (1842490 to W.A.F.).

## AUTHOR CONTRIBUTIONS

Conceptualization: M.G., C.F., K.A.J.

Data curation: R.B.H., W.A.F., C.J.F., K.A.J.

Formal analysis: R.B.H., M.G., K.A.J.

Funding acquisition: C.F., K.A.J.

Investigation: R.B.H., M.G., W.A.F., C.J.F., K.A.J.

Methodology: R.B.H., M.G., W.A.F., C.F., K.A.J.

Project administration: C.F., K.A.J.

Resources: N.L., S.A.V., L.W., R.K.P., H.Z., S.L.S.

Software: M.G. Supervision: C.F., K.A.J.

Validation: R.B.H., A.C. Visualization: R.B.H., K.A.J.

Writing – original draft: R.B.H., K.A.J.

Writing – review & editing: R.B.H., K.A.J.

---

**Figure S1.**
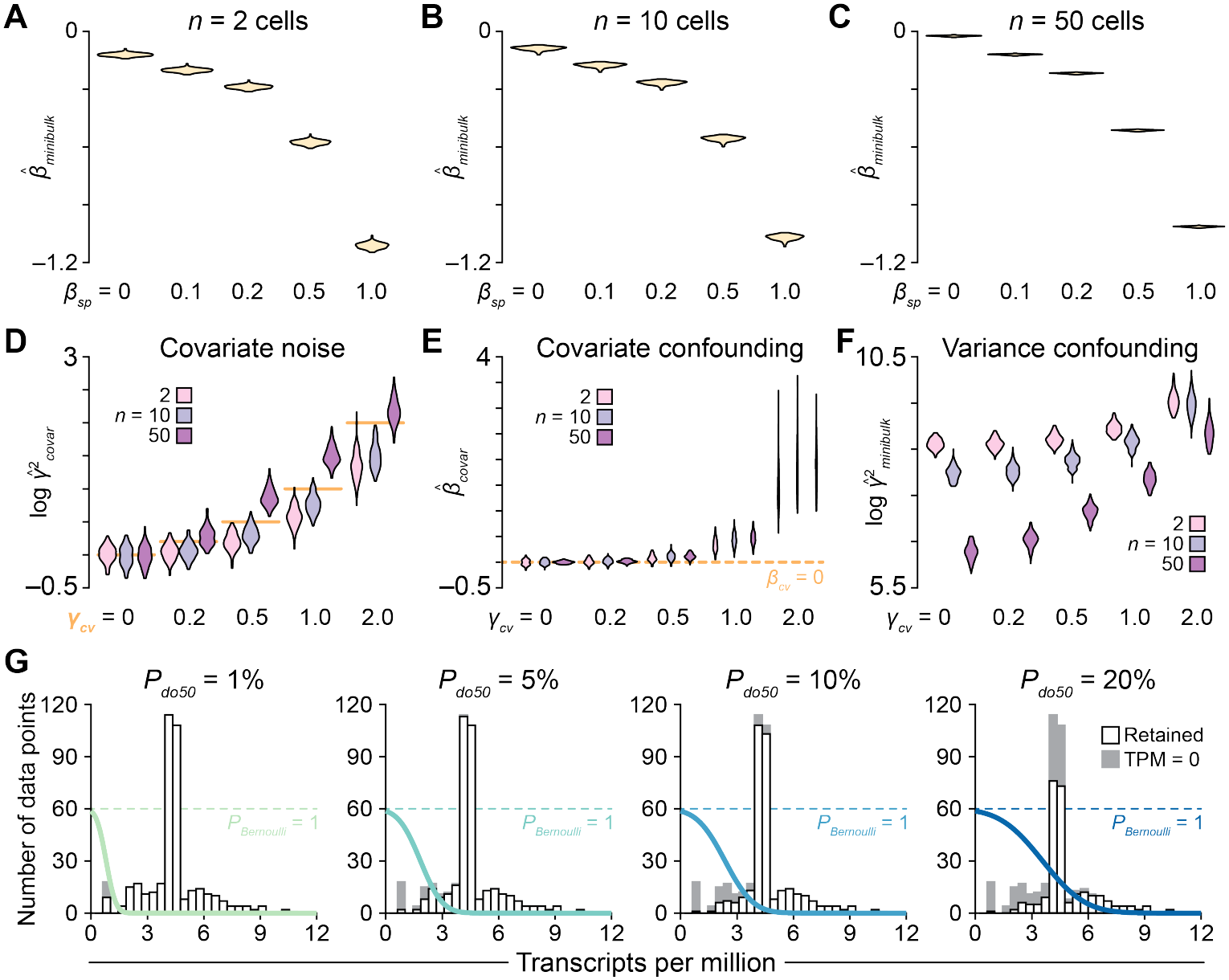
Compensatory changes and nuisance parameters in stochprofIC. Related to Figure 2. (A–C) *β^ _minibulk_* offsets the gains in *β^* _0_ caused by *β_sp_*. *β_sp_* varies where indicated for *n* = 2 cells (A), 10 cells (B), and 50 cells (C). *γ_sp_* = 0.01, *β_cv_* = 0, *γ_cv_* = 0. (D) 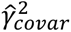 is a lagging indicator of *γ_cv_*. The orange line marks the true log 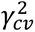 value. (E and F) Large *γ_cv_* values confound the *β^ _covar_* (E) and 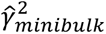 (F) estimates of stochprofIC. The orange line in (E) marks the true *β_cv_* value. (G) Illustration of the abundance-dependent Bernoulli random variable^38^ used to simulate dropouts. The Bernoulli probability decreases sigmoidally with increasing measured transcript abundance, with the midpoint defined as a percentile of the overall dataset (*P_do50_*). For (D–F), *n* and *γ_cv_* vary where indicated. *β_sp_* = 0, *γ_sp_* = 0.01, *β_cv_* = 0. For (A–F), violin plots reflect the distribution of *N* = 100 independent synthetic datasets of *k* = 100 mini-bulk samples + 100 split-pool controls x two batches.

**Figure S2.**
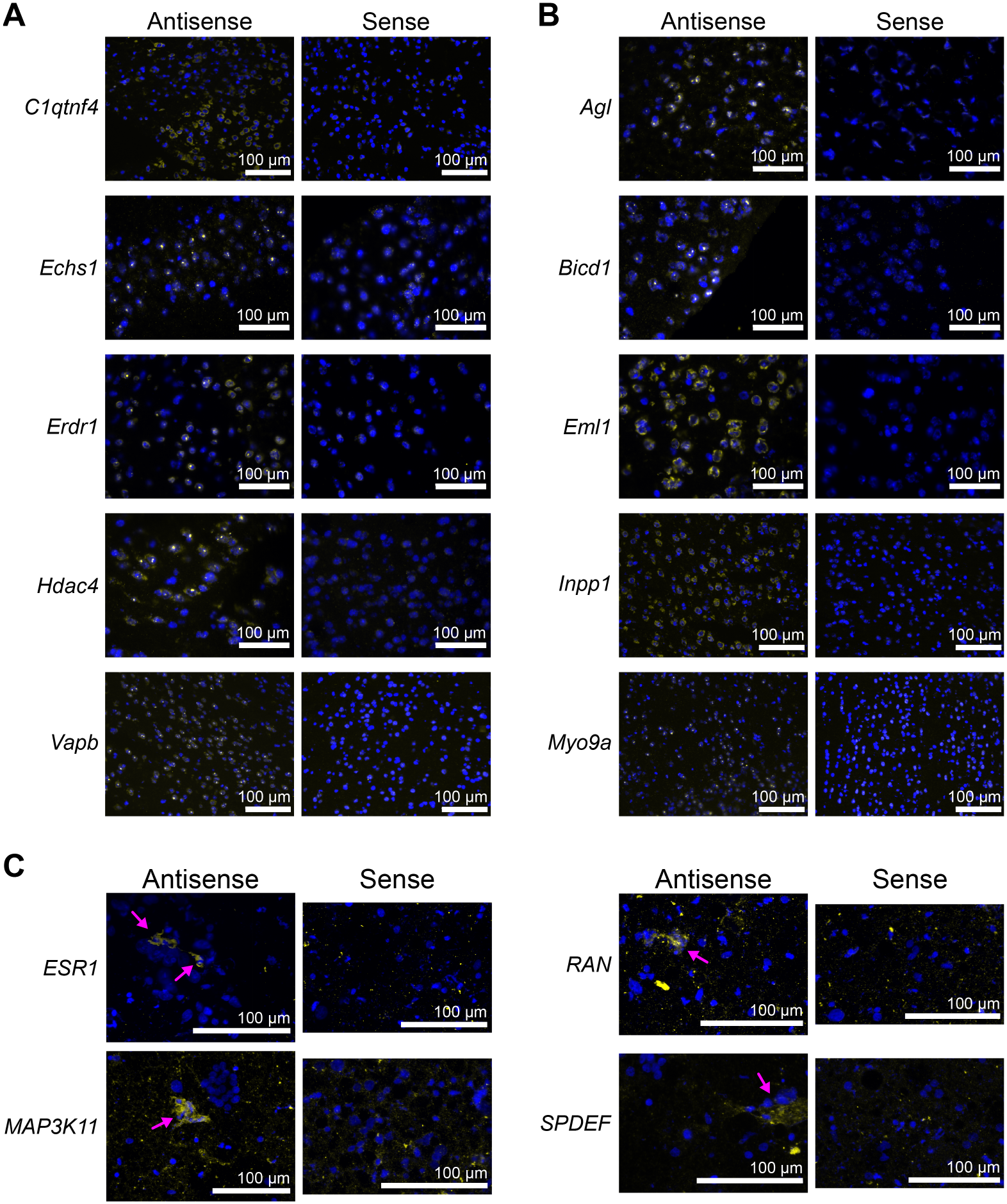
RNA FISH riboprobe validation. Related to Figures 3–6. (A and B) Antisense and sense hybridizations (yellow) of the indicated 12 dpi (A) and 90 dpi (B) gene candidates. (C) Antisense and sense hybridizations (yellow) of the indicated luminal breast cancer gene candidates. Magenta arrows indicate clusters of positive cells in the antisense images. Background fluorescence in the sense images is caused by pathology ink used to mark the tumor margins. For (A–C), the scale bar is 100 µm.

**Figure S3.**
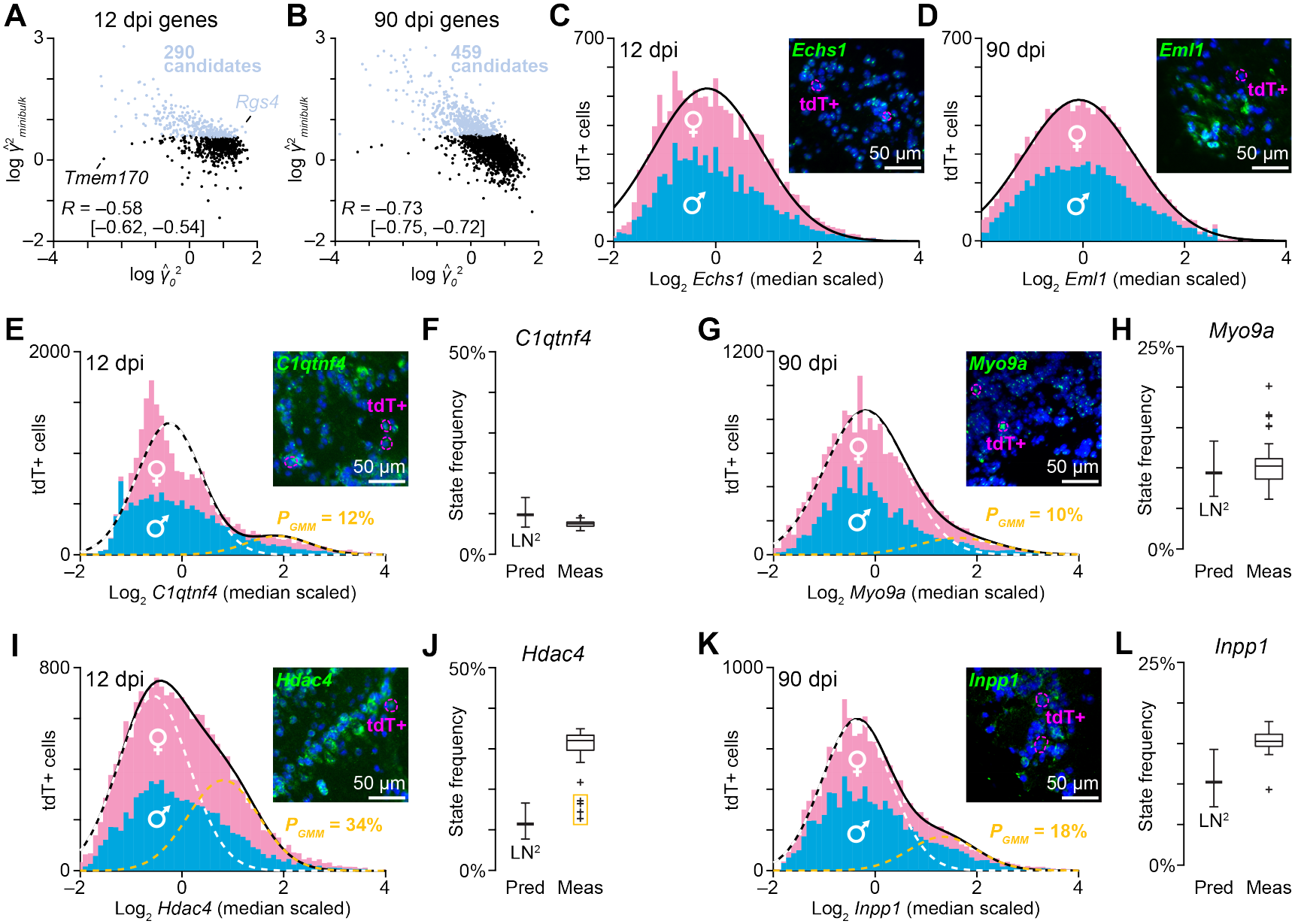
Additional stochprofIC statistics and OPC genes independently validated by RNA FISH. Related to Figure 3. (A and B) Anticorrelation between log 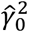 and 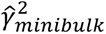 at 12 dpi (A) and 90 dpi (B). Genes deviating from the anticorrelation are annotated in (A). Genes with log *γ*^2^*_minibulk_* > 0 (*q* < 0.15) are indicated as candidates (blue). The log Pearson correlation (*R*) is shown with 90% CI in brackets. (C and D) *Echs1* at 12 dpi (C) and *Eml1* at 90 dpi (D) have one-state log-normal abundance in tdT+ OPCs from males (blue) and females (pink). (E–L) *C1qtnf4* [(E) and (F)] and *Hdac4* [(I) and (J)] at 12 dpi and *Myo9a* [(G) and (H)] and *Inpp1* [(K) and (L)] at 90 dpi have two-state log-normal abundance in tdT+ OPCs from males (blue) and females (pink). In (F), (H), (J), and (L), the measured (Meas) frequency of cells in the high expression state is compared to the predicted (Pred) frequency from stochprofML^21^ given stochprofIC-corrected data and a two-state log-normal model with equal variance (LN^2^). For (C–E), (G), (I), and (K), data are from *N* = 13504 (C), 4340 (D), 23902 (E), 6248 (G), 18249 (I), and 5152 (K) tdT+ OPCs in four mice (two males, two females) at the indicated time points. The inset shows representative RNA FISH intensities (green) in the olfactory bulb tissue with tdT+ OPCs outlined in magenta. The scale bar is 50 µm. For (F), (H), (J), and (L), predictions are shown as the maximum-likelihood estimate ± 95% CI and measurements are shown as the estimated frequency from *N* = 50 bootstrap replications of the starting total-cell population for each gene.

**Figure S4.**
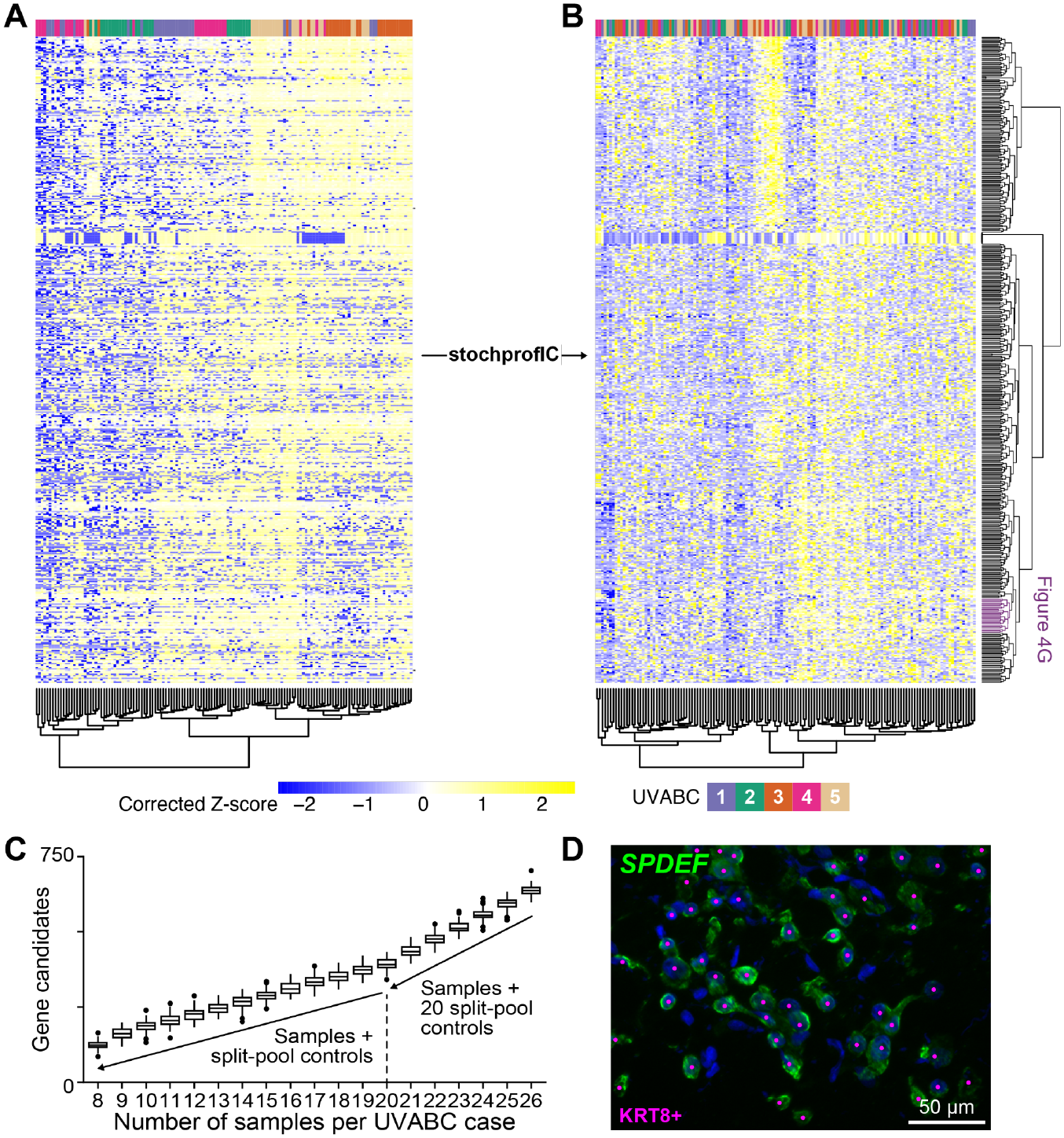
Clustering and independent validation of cell-state heterogeneities in luminal breast cancer. Related to Figure 4. (A and B) Hierarchical clustering of log-transformed, correction-eligible genes from Singh et al.^20^ with (B) or without (A) correction by stochprofIC. Row-standardized clusters in (B) were kept constant in (A), and the two datasets were separately column clustered by Euclidean distance and Ward’s linkage. Note the covariate structure in the patient track at the top of (A). The *ESR1* cluster extracted in Figure 4G is highlighted in (B, purple). (C) The number of genes with detectable log *γ*^2^*_minibulk_* > 0 (*q* < 0.15) decreases linearly with dataset size. Data from Singh et al.^20^ was repeatedly downsampled as 20–26 mini-bulk samples + 20 split-pool controls (right) or 8–20 mini-bulk samples + 8–20 split-pool controls (left) and analyzed by stochprofIC. Boxplot shows the median, interquartile range (box), 1.5 times the interquartile range (whiskers), and outliers (markers) from *N* = 100 downsampling iterations. (D) Representative RNA FISH intensities for *SPDEF* (green) in a tumor scrape (UVABC218; File S2) with KRT8+ breast cancer cells dotted in magenta. The scale bar is 50 µm.

**Figure S5.**
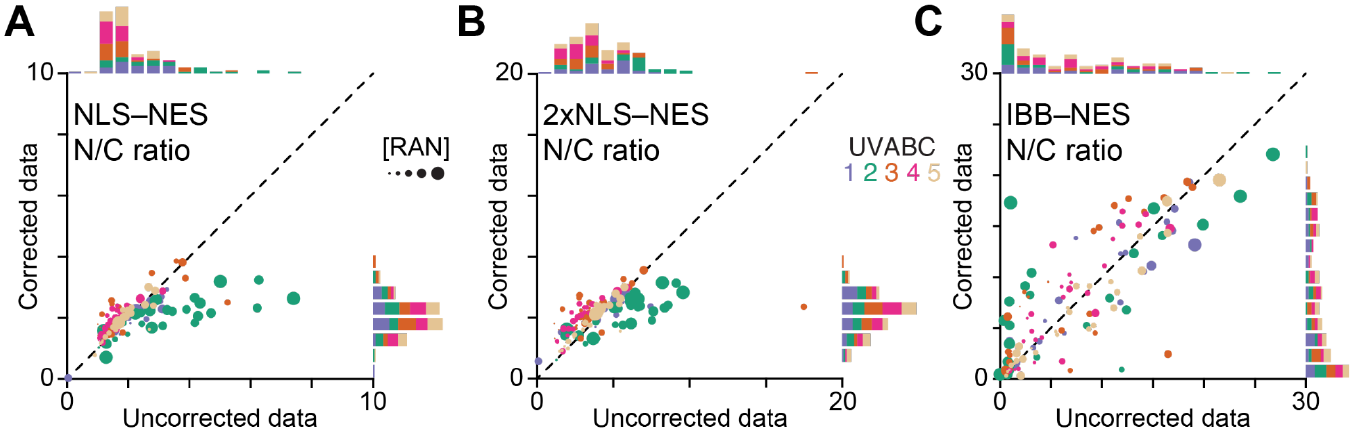
Addition of a nuclear export sequence does not influence uncorrected-vs.-corrected predictions of the nucleocytoplasmic transport model. Related to Figure 5. (A–C) Predicted steady-state nuclear-to-cytoplasmic (N/C) ratio of the indicated cargo with an appended nuclear export sequence (NES) when uncorrected initial conditions (x-axis) or stochprofIC-corrected initial conditions (y-axis) are used in the model. UVABC cases are color-coded as in Figure 5D–G, and marker size is proportional to the square root of uncorrected RAN abundance. See Wang et al.^33^ for further details on cargo and the model.

**Figure S6.**
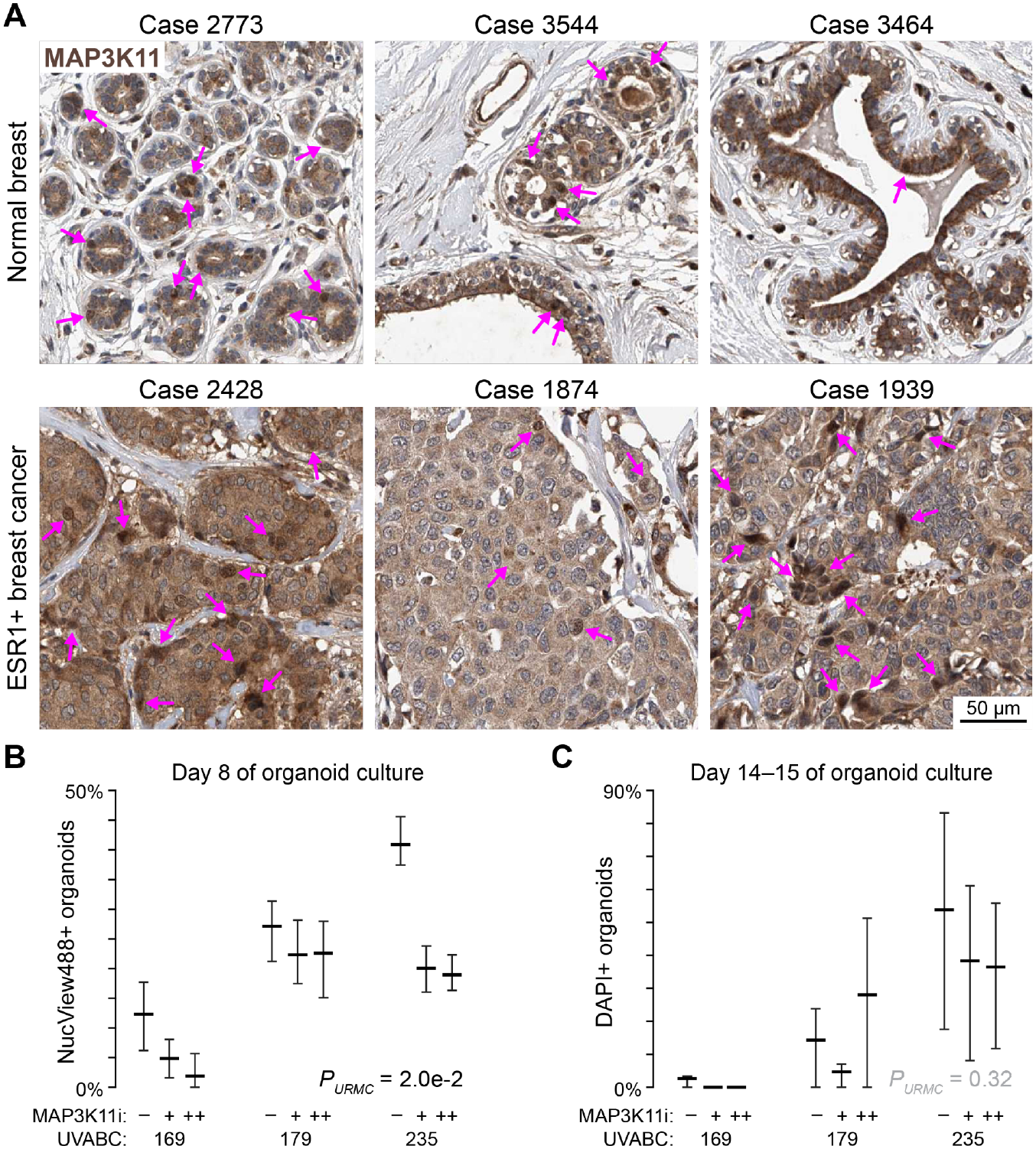
A luminal MAP3K11-high cell state linked to stress-induced cell death. Related to Figure 6. (A) Immunohistochemical stains for MAP3K11 in normal breast tissue (top) and ESR1+ breast cancer (bottom) from The Human Protein Atlas.^56^ ESR1 positivity was determined by immunohistochemistry in the paired case number. Magenta arrows indicate cells with elevated immunoreactivity. The scale bar is 50 µm. (B) Same cultures as in Figure 6E analyzed for NucView488 live-cell caspase reporter activity on day 8. (C) MAP3K11 inhibition with URMC-099 has mixed effects on overall cell death of zero-passage luminal breast cancer organoids. At the culture endpoint, cultures were assessed for membrane permeabilization with dilute DAPI (STAR Methods). Quantification of DAPI+ organoid frequencies from *N* = 3 independent patients. For (B) and (C), data are shown as the mean ± 84% bootstrapped confidence intervals,^70^ and the fixed effect of URMC-099 (*P_URMC_*) was assessed as a continuous factor by multiway ANOVA of arcsine square-root transformed percentages with UVABC case as an additional fixed effect.

## STAR METHODS

### KEY RESOURCES TABLE

| **REAGENT or RESOURCE** | **SOURCE** | **IDENTIFIER** |
|---|---|---|
| **Antibodies** |  |  |
| Alexa Fluor 594 goat anti-mouse | Invitrogen | Cat# A11005 |
| Alexa Fluor 594 goat anti-rabbit | Invitrogen | Cat# A11012 |
| Alexa Fluor 647 donkey anti-sheep | Invitrogen | Cat# A21448 |
| Living Colors rabbit anti-dsRed | Takara | Cat# 632496 |
| Mouse anti-cytokeratin 8 | Sigma Aldrich | Cat# MAB3414 |
| Rabbit anti-cytokeratin 8 | Abcam | Cat# ab53280 |
| Sheep anti-digoxigenin | Millipore Sigma | Cat# 11333089001 |
| **Bacterial and virus strains** |  |  |
| Stellar Competent Cells | Takara | Cat# 636766 |
| **Biological samples** |  |  |
| *Cspg4-CreER; Rosa-tdT; Nf1<sup>F/F</sup>; p53<sup>F/F</sup>* mouse olfactory bulbs | Sutcliffe et al.<sup>29</sup> | N/A |
| Human ER+ breast cancer scrapes | This paper | File S2 |
| **Chemicals, peptides, and recombinant proteins** |  |  |
| 0.1 M DTT | Invitrogen | Cat# Y00147 |
| 20x SSC | Promega | Cat# V4261 |
| 5x First Strand Buffer | Invitrogen | Cat# Y02321 |
| 5x Phusion GC Reaction Buffer | NEB | Cat# B0519S |
| 5x Phusion HF Reaction Buffer | NEB | Cat# B0518S |
| A83-01 | Millipore Sigma | Cat# SML0788 |
| Acetic anhydride | Millipore Sigma | Cat# 242845 |
| adDMEM/F12 | Thermo Fisher | Cat# 12634-028 |
| Animal-Free Recombinant Human EGF | Peprotech | Cat# AF-100-15 |
| *BamHI* HF | NEB | Cat# R3136L |
| B-27 Plus supplement (50x) | Thermo Fisher | Cat# A3582801 |
| Betaine | Sigma | Cat# B0300 |
| β-Estradiol | Sigma-Aldrich | Cat# E2257 |
| Bovine Serum Albumin | Millipore Sigma | Cat# A600-5G |
| Calf intestinal alkaline phosphatase | Invitrogen | Cat# 18009-027 |
| Collagenase Type II | Thermo Fisher | Cat# 17101015 |
| Dextran sulfate, 50% | Chemicon | Cat# 3730 |
| *EcoRI* HF | NEB | Cat# R3101L |
| Ethanol, 200 proof | Decon | Cat# 2716 |
| FGF-7 | Peprotech | Cat# AF-100-19 |
| FGF-10 | Peprotech | Cat# AF-100-26 |
| Formamide | Millipore Sigma | Cat# 47671 |
| Forskolin | Millipore Sigma | Cat# F6886 |
| GlutaMAX | Thermo Fisher | Cat# 35050-061 |
| Glycogen | Invitrogen | Cat# 10814010 |
| Hepes | Gibco | Cat# 15630080 |
| Hydrochloric acid, 12 N | Fisher | Cat# A481-212 |
| Hydrocortisone | Sigma-Aldrich | Cat# H0888 |
| Matrigel | Corning | Cat# 356253 |
| N-acetyl-L-cysteine | Sigma-Aldrich | Cat# A9165 |
| Neg-50 | Epredia | Cat# 6502 |
| Nicotinamide | Millipore Sigma | Cat# NO636 |
| NucView 488 Green Caspase-3 Dye (DMSO) | Millipore Sigma | Cat# SCT100 |
| NucView 488 Green Caspase-3 Dye (PBS) | Millipore Sigma | Cat# SCT101 |
| Paraformaldehyde | Fisher | Cat# 41678-5000 |
| Penicillin-Streptomycin | Thermo Fisher | Cat# 15140-122 |
| Phenol/chloroform/isoamyl alcohol (25:24:1 mixture; pH 6.7/ 8.0) | Fisher | Cat# BP1752I-100 |
| Phusion HF DNA Polymerase | NEB | Cat# M0530L |
| Primocin | Invivogen | Cat# Ant-pm-1 |
| rCutSmart Buffer | NEB | Cat# B6004S |
| Recovery Cell Freezing Medium | Gibco | Cat# 12648010 |
| Red blood cell lysis buffer | Millipore Sigma | Cat# 11814389001 |
| SB 202190 | Sigma-Aldrich | Cat# S7067 |
| Superscript III reverse transcriptase | Invitrogen | Cat# 56575 |
| T4 DNA Ligase | NEB | Cat# M0202S |
| T4 DNA Ligase Buffer | NEB | Cat# B0202S |
| Triethanolamine | Alfa Aesar | Cat# L04486 |
| Tween-20 | Fisher | Cat# BP337-500 |
| URMC-099 | MedChemExpress | Cat# HY-12599 |
| Western Blocking Reagent | Roche | Cat# 11921673001 |
| Y-27632 dihydrochloride | StemCell | Cat# 72304 |
| Yeast tRNAs | Invitrogen | Cat# 15401-029 |
| **Critical commercial assays** |  |  |
| Direct-zol RNA Miniprep Kit | Zymo Research | Cat# R2052 |
| MAXIscript Transcription Kit | Invitrogen | Cat# AM1322 |
| PureLink Quick Gel Extraction Kit | Invitrogen | Cat# K2100-25 |
| Qubit RNA BR Assay Kit | Invitrogen | Cat# Q10210 |
| TURBO DNase Kit | Ambion | Cat# AM1907 |
| **Deposited data** |  |  |
| Breast cancer single-cell atlas | Wu et al.<sup>6</sup> | GSE176078 |
| Human Protein Atlas | Uhlén et al.<sup>56</sup> | Normal: 65747_B_1_4, 65747_B_2_4, 65747_B_3_4<br>Breast cancer: 66954_A_5_3, 66954_A_6_1, 66954_A_6_5 |
| Normal breast tissue single-cell atlas | Kumar et al.<sup>54</sup> | GSE235326 |
| **Experimental models: Cell lines** |  |  |
| Noggin-producing 293T cells | Clevers group, Hubrecht Institute | N/A |
| R-Spondin-producing 293T cells | R&D Systems | Cat# 3710-001-01 |
| Wnt3a-producing L cells | Clevers group, Hubrecht Institute | N/A |
| **Experimental models: Organisms/strains** |  |  |
| **Oligonucleotides** |  |  |
| PCR cloning primers for RNA FISH riboprobes | This paper | File S3 |
| **Recombinant DNA** |  |  |
| pcDNA3 Agl.p | This paper | Addgene #265048 RRID: Addgene_265048 |
| pcDNA3 Bicd1.p | This paper | Addgene #265051 RRID: Addgene_265051 |
| pcDNA3 C1qtnf4.p | This paper | Addgene #265043 RRID: Addgene_265043 |
| pcDNA3 Echs1.p | This paper | Addgene #265046 RRID: Addgene_265046 |
| pcDNA3 Eml1.p | This paper | Addgene #265052 RRID: Addgene_265052 |
| pcDNA3 Erdr1.p | This paper | Addgene #265047 RRID: Addgene_265047 |
| pcDNA3 ESR1.p | This paper | Addgene #265053 RRID: Addgene_265053 |
| pcDNA3 Hdac4.p | This paper | Addgene #265044 RRID: Addgene_265044 |
| pcDNA3 Inpp1.p | This paper | Addgene #265049 RRID: Addgene_265049 |
| pcDNA3 MAP3K11.p | This paper | Addgene #265054 RRID: Addgene_265054 |
| pcDNA3 Myo9a.p | This paper | Addgene #265050 RRID: Addgene_265050 |
| pcDNA3 RAN.p | This paper | Addgene #265055 RRID: Addgene_265055 |
| pcDNA3 SPDEF.p | This paper | Addgene #265056 RRID: Addgene_265056 |
| pcDNA3 Vapb.p | This paper | Addgene #265045 RRID: Addgene_265045 |
| **Software and algorithms** |  |  |
| CellProfiler (version 4.2.8) | Stirling et al.<sup>71</sup> | https://cellprofiler.org/releases |
| gamlss (version 5.5) | CRAN | https://cran.r-project.org/web/packages/gamlss/ |
| MATLAB (version R2026a) | Mathworks | https://www.mathworks.com/downloads/ |
| OrganoSeg2 | Wells et al.<sup>72</sup> | https://github.com/JanesLab/OrganoSeg2 |
| Pheatmap (version 1.0.13) | CRAN | https://cran.r-project.org/web/packages/pheatmap/ |
| stochprofIC | This paper | https://github.com/fuchslab/stochprofIC |
| stochprofML (version 2.0.3) | Bajikar et al.<sup>21</sup> | https://github.com/fuchslab/stochprofML/ |
| umap (version 0.2.10.0) | CRAN | https://cran.r-project.org/web/packages/umap/ |
| **Other** |  |  |
| 22 mm x 50 mm cover glass | Electron Microscopy Sciences | Cat# 72204-04 |
| 22 mm x 22 mm cover glass | Fisher | Cat# 12-541-B |
| 96-well (1 x 8 Stripwell™) Clear Flat Bottom Polystyrene TC-treated Microplates | Corning | Cat# 9102 |
| CFX96 Real-Time PCR System | Biorad | Cat# 3600037 |
| Cell strainer, 100 µm | Avantar | Cat# 77972-876 |
| CryoStar NX50 cryostat | Epredia | Cat# 95-710-0L |
| EVOS M7000 slide scanner | Thermo Fisher | Cat# AMF7000 |
| Nuclease-free water | Ambion | Cat# AM9937 |
| SuperFrost Plus Slides | Avantor | Cat# 48311-703 |
| Tissue-Tek Cryomold | Sakura | Cat# 4566 |

### RESOURCE AVAILABILITY

#### Lead Contact

Further information and requests for resources and reagents should be directed to and will be fulfilled by the Lead Contact, Kevin A. Janes.

#### Materials Availability

Plasmids generated in this study have been deposited at Addgene (#265043–#265056).

#### Data and Code Availability

- Source data will be deposited at LabArchives and be publicly available as of the date of publication. The DOI will listed in the key resources table.
- All original code has been deposited at GitHub and is publicly available as of the date of publication. Scripts used to generate figures will be deposited at LabArchives and be publicly available as of the date of publication. DOIs will be listed in the key resources table.
- Any additional information required to reanalyze the data reported in this paper is available from the Lead Contact upon request.

### EXPERIMENTAL MODEL AND STUDY PARTICIPANT DETAILS

#### Mice

Cryobanked olfactory bulb tissue of unused *Cspg4-CreER; Rosa-tdT; Nf1^F/F^; p53^F/F^* mice (mixed background) from Sutcliffe et al.^29^ was freshly sectioned. In these animals, tamoxifen dosing began at P45 for five days as described.^29^ We selected samples from two males and two females each at 12 dpi and 100 (∼90) dpi for this study. Time points were randomized by cage. Before sacrifice, mice were housed together on a 10-hour dark cycle (8 pm–6 am) and 14-hour light cycle (6 am–8 pm) at 21.5 °C and 31.5% relative humidity. All work with animals was performed under Protocol #3955 in accordance with the guidelines and regulations of the National Institutes of Health and the Institutional Animal Care and Use Committee of the University of Virginia, which is accredited by the Association for the Assessment and Accreditation of Laboratory Animal Care.

#### Patients

Primary breast cancer scrapes were obtained as remnant medical waste from women undergoing surgery at the University of Virginia Health System. Age, self-reported race and ethnicity, and other clinical characteristics are reported in File S2 along with their usage in the study. Sample acquisition and experimental procedures were performed by waiver of consent under 45CFR46.116 of the 2018 Common Rule and Protocol #14176 approved by the Institutional Review Board for Health Sciences Research at the University of Virginia.

### METHOD DETAILS

#### Synthetic Mini-bulk Datasets

Technical noise and covariate parameters were evaluated atop a previously used^21^ two-state distribution: *μ*_1_ = 0.5, *μ*_2_ = –2.5, *σ* = 0.225, and *F* = 22.5%. Binomial and lognormal pseudorandom numbers were used to generate the mini-bulk mixture of *n* cells as before.^21^ Each evaluation comprised *k* observations from the mini-bulk distribution that were then multiplied with pseudorandom lognormal numbers for technical noise (*LN*(*β_sp_*, *γ_sp_*)) and covariate effects for the second batch relative to the first (*LN*(*β_cv_, γ_cv_*)). Split-pool samples were generated by taking the mean of all mini-bulk observations for an evaluation and multiplying by technical noise. Parameters for the reference case were *β_sp_* = 0.01, *γ_sp_* = 0.2, *β_cv_* = 0.4, and *γ*_cv_ = 0.02. Noise and covariate parameters were perturbed independently while keeping all other parameters fixed. For each evaluation, simulations were replicated 100 times to estimate parameter uncertainties related to sampling.

Where indicated, we overlaid dropouts on synthetic datasets using a probabilistic model.^38^ The stochastic dropout of observation *i* (*D_i_*) given its expression value (*z_i_*) was modeled as:

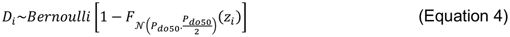

where 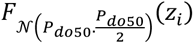 is the cumulative distribution function for a normal distribution with mean of *P_do50_* (defined as a percentile of the synthetic data in a simulation) and standard deviation of *P_do50_*/2. If the Bernoulli outcome was true, *z_i_* was changed to zero; if false, *z_i_* was retained.

#### StochprofIC

The package takes as input a tibble with columns corresponding to sample name, gene name, measurement type (split-pool control or mini-bulk sample), categorical fixed effect (e.g., sex, patient), and abundance (e.g., TPM). The function stochprofIC.screening evaluates genes for robust dispersion by trimming the three highest and lowest values (as implemented^20^ in Pagoda), converting remaining zeros to NaN, and then fitting to the normal model in Equation 1 using the gamlss package in R. (Note that the stochprofIC implementation of gamlss performs distributional regression on location and scale, but not shape.) Estimates of log 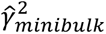 are screened at a Šidák-corrected familywise error rate of 5% before the testing phase. The function stochprofIC.testing takes the screened genes, adds one to the TPM values, and then fits to the normal model in Equation 1 after log transformation. In the testing step, no trimming or NaN conversion is performed. Genes with log *γ*^2^*_minibulk_* > 0 at a false-discovery rate of 15% were deemed candidates for correction, which involved subtracting estimated log means and dividing by log variances as in Equation 3.

#### Pagoda

Sutcliffe et al.^29^ adopted a leave-one-out crossvalidation approach to defining overdisperse genes, which was not directly comparable with stochprofIC. Therefore, we repeated the same analytical pipeline on eligible transcripts^20^ using all the mini-bulk and split-pool observations for i) 12 dpi males, ii) 12 dpi females, iii) 90 dpi males, and iv) 90 dpi females. Adjusted variance results for these four separate analyses are included in File S1.

#### Cloning

Riboprobes were PCR cloned into pcDNA3 as 150–225 bp fragments with *EcoRI* and *BamHI* restriction sites on the 5’ and 3’ ends respectively (File S3). The PCR template was a cDNA library prepared by oligo(dT)_24_-primed reverse transcription of TURBO DNAse-treated RNA as described.^73^ RNA was extracted and pooled from four mouse brains (two male + two female) that were stored in TRIzol at – 80°C, thawed–homogenized, and purified using the Direct-zol RNA mini-prep kit (Zymo). All cloned riboprobes were verified by restriction digest and Sanger sequencing and are deposited with Addgene (#265043–#265056).

#### RNA Fluorescence in situ hybridization (FISH)-Immunofluorescence

Starting with an existing RNA FISH protocol,^26^ we added a paired immunofluorescence step that co-stained for tdT (mouse brain) or KRT8 (breast cancer) alongside riboprobe haptens after hybridization.

##### Riboprobe synthesis

Digoxin-labeled riboprobes were prepared by *in vitro* transcription as described.^26^ Riboprobe concentrations were estimated using the Qubit RNA BR assay (Thermo Fisher) on a CFX96 Real-Time PCR System (Bio-Rad), and stocks were stored at –80°C.

##### Mouse brain cryosections

Olfactory bulb tissue from *Cspg4-CreER; Rosa-tdT; Nf1^F/F^; p53^F/F^* animals was freshly cryoembedded in Neg-50 (Epredia) and stored at –80°C before cutting 8 µm sections at –40°C on a CryoStar NX50 cryostat (Epredia). Cryosections (two per slide) were rapidly refrozen after wicking and stored at –80°C before use.

##### Breast cancer scrapes

Scrapes were obtained after gross dissection of the tumor mass and normal pathological accession. A clean razor blade was scraped along the tumor mass, and scraped cells were tapped and smeared onto SuperFrost Plus microscope slides (Avantor). Slides were kept on ice in a 50 ml conical tube for transport until fixation in 3.7% paraformaldehyde for 15 minutes at room temperature, followed by serial dehydration in 70% ethanol for two minutes and 95% ethanol for two minutes before storage in 100% ethanol at –20°C until use.

##### Hybridization and immunostaining

Slides were brought to room temperature, acid denatured, postfixed, acetylated, dehydrated, and hybridized overnight with 4 ng/µl riboprobe as described.^26^ After washing, slides were blocked as before and costained for one hour at room temperature with antibodies recognizing digoxin (Millipore; sheep, 1:200 dilution) and tdT (Takara; rabbit, 1:100 dilution) or KRT8 (Millipore; mouse, 1:100 dilution or Abcam; rabbit, 1:100 dilution). Slides were washed three times with PBS for five minutes each, incubated with AlexaFluor-conjugated anti-species secondary antibodies, washed, counterstained, and mounted as described.^26^

#### Image Acquisition

Slides were imaged on EVOS M7000 slide scanner (Invitrogen) with a UPlanSApo 20x 0.85 NA oil immersion objective (Olympus). Slides were illumination, exposure, and gain matched for each target with the following LED light cubes: DAPI (excitation: 357 nm; emission: 417–477 nm), Texas Red (excitation: 585 nm; emission: 612–644 nm), and Cy5 (excitation: 635 nm; emission: 672–712 nm). Images were acquired as tiled composites for an entire olfactory bulb section (35–88 images) or for a representative section of a tumor scrape (165–352 images, depending on cellularity). Two serial olfactory bulb sections were imaged per riboprobe per mouse, and one tumor scrape was imaged per riboprobe per case.

#### Systems Modeling of Nucleocytoplasmic Transport

We used MODEL2210060001 to predict steady-state nucleocytoplasmic localization of different cargo as previously described.^33^ To parameterize initial conditions, we extracted uncorrected and stochprofIC-corrected TPM data from UVABC mini-bulk samples for the following genes: *RAN*, *RANBP1*, *RANGAP1*, *RCC1*, *NUTF2* (common name: NTF2), all *KPNA* isoforms excluding pseudogenes, *KPNB1*, *CSE1L*, *XPO1* (common name: CRM1), *RANBP3*, all *NUP*-prefixed subunits of the nuclear pore complex. NUP subunits were median scaled to reflect their fixed stoichiometry in the nuclear pore complex, and subunits with a median of zero were removed. Importin-α was estimated by summing TPM values of *KPNA* isoforms, and nuclear pore complexes were estimated by summing the median-scaled NUP subunits. Remaining uncorrected observations with zero TPM were taken as not detected and assigned the median non-zero TPM for that species. Uncorrected and stochprofIC-corrected species were scaled to their respective medians, and the square root of these median-normalized values was used to scale model initial conditions previously collected in a breast epithelial line.^33^ The square-root transformation was implemented to reflect nonlinear RNA-protein relationships^65^ and attenuate the spread of initial conditions among mini-bulk samples. As before,^33^ initial conditions were run to steady state followed by a step increase in 1 µM cargo containing different combinations of import and export sequences. The steady-state abundances of nuclear and cytoplasmic cargo were ratioed and taken as the final output of each model.

#### Organoids

Tumor scrapes were isolated from glass slides, digested, and plated immediately for organoid culture as described^58^ or cryopreserved in Recovery Cell Freezing Medium (Gibco). Frozen cells (UVABCO169 and UVABCO179) were rapidly thawed from liquid nitrogen storage and plated immediately for organoid culture. Organoids were maintained as before,^58^ and URMC-099 at the indicated concentrations plus 0.1% DMSO was added to the day 4 medium change and maintained for the rest of the culture. Cell death imaging with NucView 488 Green Caspase-3 Dye (Sigma) added at day 4 and DAPI at the day 14 or day 15 endpoint was performed as described.^72^

### QUANTIFICATION AND STATISTICAL ANALYSIS

#### Uniform Manifold Approximation and Projection (UMAP)

TPM data from GSE147356 were log_2_(TPM+1) transformed as previously described.^20^ Endogenous transcripts for mini-bulk samples, split-pool controls, patient-specific medians (calculated across all mini-bulks and split-pools) were projected with the umap package in R and the following non-default parameters: n_neighbors = 4, metric = “pearson”, and min_dist = 0.0001.

#### StochprofML

Single-cell distributions were estimated from stochprofIC-corrected mini-bulk data by using stochprofML in R to compare LN^2^ and rLN^2^ best-fit models to one-state LN by BICw. The package also estimates a mixture of one exponential distribution (representing a silent expression state) and one log-normal distribution (EXP–LN). However, this mixture was deemed impractical for validation by RNA FISH, because autofluorescence, nonspecific probe binding, and detector shot noise blur the silent expression state into a second log-normal distribution. 95% CIs for expression frequency were estimated by computing the inverse Hessian of the negative log-likelihood at the best-fit parameters as described.^21^ If the Hessian could not be computed for certain rLN^2^ models, best-fit *µ*_1_, *µ*_2_, and *σ* parameters for each gene were used to examine possible *F* values using the stochprofML function ‘stochprof.search.rLNLN’. Parameter value ranges were set tightly (ranges of 0.01 to 1) around each best-fit value while *F* was left unconstrained. A grid search was performed 10,000 times for each gene to find a minimum negative-log-likelihood proportion value. Minimum negative log-likelihood plus half the 95^th^ percentile chi-square distribution (df = 1) established thresholds for the empirical 95% confidence interval.

#### Digital Image Segmentation and Analysis

Images were segmented with CellProfiler.^71^ Cell nuclei were identified by suppressing DAPI objects under 20 pixels, gating objects to within 20–75 pixels (6.2–23 µm) in diameter, and filtering for nuclei within 500–2500 pixels^2^ (47–237 µm^2^) in area. Nuclear masks were expanded by 5 pixels (1.5 µm) and the 75th percentile of fluorescence intensity was measured for the whole mask in each channel.

Cell intensities transcripts and protein markers (tdT or KRT8) were scaled by the median intensity taken across all cells for a mouse or patient. Then, a global tdT or KRT8 threshold intensity was set at the 85th percentile of all scaled cells across mice or patients in a study. Marker-positive cells were next log-centered by the geometric mean and log-scaled by the log standard deviation calculated by sex (for OPCs) or by patient (for breast cancer cells). This re-scaling is equivalent to the harmonization of stochprofIC. Re-scaled marker-positive cells were aggregated and fit to a log-normal distribution or a mixture of two log-normal distributions with equal or unequal variance depending on the best-fit model of stochprofML according to BICw. Bootstrapping of the entire procedure was performed by resampling the original cell intensities with replacement 50 times using a seed that ensured 100% successful log-normal fits. Representative image fields were selected based on marker positivity and K-S distance to the overall marker-positive population.

For zero-passage organoids, images were segmented with OrganoSeg2 by adjusting segmentation parameters for each case (File S4). Brightfield segments were overlaid on fluorescence images of NucView488 and DAPI to define organoid boundaries. The fluorescence of each segment was summarized by its 95^th^ percentile intensity except for UVABC235, which contained dead-cell debris and thus was summarized by the 70^th^ percentile for DAPI as before.^72^ Summarized intensities were pooled for all timepoints in each case and centered by subtracting the aggregate 20th percentile intensity. Centered values were then aggregated for all cases, and a single Gaussian was built in MATLAB using nlinfit and constant error. We used the 95^th^ percentile (NucView488) or the 90^th^ percentile (DAPI) of the Gaussian fit to score organoids as positive or negative. An organoid that exceeded the NucView488 threshold was considered apoptotic at every timepoint thereafter.

#### Hierarchical Clustering

Clustering was performed after row standardization with the pheatmap function in R or the clustergram function in MATLAB using a Euclidean distance metric and Ward’s linkage.

#### Statistics

Contingency table enrichments were evaluated by the hypergeometric test, and off-diagonal biases were evaluated by the sign-rank test. Uncertainty in organoid percentages was estimated by bootstrapping organoid segments with replacement 100 times for each case, day, and inhibitor concentration. Intensities from each iteration of bootstrapping were aggregated, recentered and refit as above to calculate the cutoff for each bootstrap replicate, and the resulting values were summarized as an estimated 84% CI.^70^ Percentages on a given day of organoid culture were arcsine square-root transformed and analyzed by two-way ANOVA in MATLAB using anovan without interaction, with inhibitor dose as a continuous fixed effect, and with UVABCO case as a categorical fixed effect.

## SUPPLEMENTAL FILE TITLES AND LEGENDS

**File S1. Gene lists for the glioma premalignancy study. Related to Figure 3**. This file contains eight sheets: Pagoda overdispersion analysis results from the complete Sutcliffe et al.^29^ dataset for i) males at 12 dpi, ii) females at 12 dpi, iii) males at 90 dpi, and iv) females at 90 dpi; stochprofIC estimates of *β^*_0_, 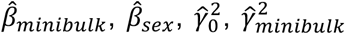, and 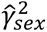 at v) 12 dpi and vi) 90 dpi; and Bayesian Information Criterion weights (BICw) and two-state expression frequencies (p1) for competing stochprofML models at vii) 12 dpi and viii) 90 dpi.

**File S2. Gene lists for the luminal breast cancer study. Related to Figure 4**. This file contains three sheets: i) stochprofIC estimates of 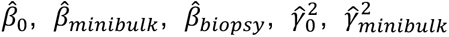, and 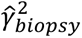; ii) Bayesian Information Criterion weights (BICw) and two-state expression frequencies (p1) for competing stochprofML models; and iii) a table of clinical characteristics for the tumor scrapes used in this study.

**File S3. Oligonucleotide sequences used for PCR cloning of RNA FISH riboprobes. Related to Figures 3–6**. This file contains one sheet of forward and reverse sequences used to clone into the *BamHI* and *EcoRI* sites of pcDNA3 along with the verified sense probe sequence.

**File S4. OrganoSeg2 segmentation parameters. Related to Figure 6**. This file contains one sheet with the segmentation settings used for each UVABC case.

